# Impact of Acute Endurance Exercise on Alternative Splicing in Skeletal Muscle

**DOI:** 10.1101/2024.11.21.624690

**Authors:** Alexander Ahn, Jeongjin J. Kim, Aaron L. Slusher, Jeffrey Y. Ying, Eric Y. Zhang, Andrew T. Ludlow

**Author notes:** Corresponding author: Andrew T. Ludlow, PhD, Ann Arbor, Michigan, USA, 830 North University Avenue, Ann Arbor, Michigan 48109-1048, USA.

## Abstract

II:

**Purpose:** Alternative splicing (AS) is a highly conserved post-transcriptional mechanism, generating mRNA variants to diversify the proteome. Acute endurance exercise appears to transiently perturb AS in skeletal muscle, but transcriptome-wide responses are not well-defined. We aimed to better understand differential AS (DAS) and differential isoform expression (DIE) in skeletal muscle by comparing short-read (SRS) and long-read RNA sequencing (LRS) data.

**Methods:** Publicly accessible SRS of clinical exercise studies were extracted from the Gene Expression Omnibus. Oxford Nanopore LRS was performed on mouse gastrocnemius before and following treadmill exercise (30m running, *n*=5 mice/group, 20 total, 10 weeks old). Differential gene expression (DGE) and DIE were analyzed and validated using RT-PCR and immunoblots.

**Results:** Both SRS and LRS illustrated significant DGE in skeletal muscle post-exercise, including 89 RNA-binding proteins (RBPs). rMATS analysis of SRS revealed that exon-skipping and intron-retaining events were the most common. Swan analysis of LRS revealed several common genes across post-exercise cohorts with significant DAS but no DGE: 13 exercise-associated genes, including *mSirt2* (24.5% shift at 24hr post-exercise [24pe], *p*=0.005); 61 RBPs, including *mHnrnpa3* (28.5% at 24pe, *p*=0.02), *mHnrnpa1* (30.6% at 24pe, *p*=0.004), and *mTia1* (53.6% at 24pe, *p*=0.004).

**Conclusions:** We illustrated that acute endurance exercise can elicit changes in AS-related responses and RBP expression in skeletal muscle, especially at 24pe. SRS is a powerful tool for analyzing DGE but lacks isoform detection, posing a major gap in knowledge of “hidden” genes with no transcriptional but significant DIE and protein expression changes. Additionally, LRS can uncover previously unknown transcript diversity and mechanisms influencing endurance exercise adaptations and responses.

## 1 INTRODUCTION

The positive impact of exercise on one’s health is undeniable. With skeletal muscle accounting for approximately 40% of body mass in adults, it is the most directly affected tissue from both the beneficial effects of physical activity as well as the deleterious effects of sedentarism. However, a large portion of the population fails to meet the minimum amount of exercise to reduce health risks. The U.S. Centers for Disease Control and Prevention reported overall prevalence of physical inactivity was 25.3% in 2020,^1^ while causing 9% of premature death globally.^2^ Why people fail to meet the minimum amount of exercise are varied but some are simply unable to even if desired: for example, muscle dystrophy, motoneuron-related sclerosis diseases, and obesity-induced osteoarthritis. To fully harness the health-promoting power of endurance exercise, we must continue to uncover the fundamental mechanisms of molecular responses and adaptations elicited by the exercise stimulus.

Endurance exercise promotes disease-free longevity via the enhancement of skeletal muscle oxidative capacity among other important responses and adaptations.^3^ Exercise is a potent cellular stressor (e.g., mechanical, metabolic, oxidative stress, etc.) that triggers a series of responses and adaptations.^4–7^ Part of the adaptive process in skeletal muscle is changes in gene expression regulation that regulates the protein landscape and alters structural and functional outcomes. Presently, reports have focused on changes in transcript abundance and differential gene expression (DGE).^8,9^ For example, endurance exercise training increases mitochondrial density, muscle-specific angiogenesis, and improves calcium handling in slow-twitch skeletal myofibers by altering the expression of the genes encoding peroxisome proliferator-activated receptor gamma coactivator 1 alpha (*PPARGC1A*, PGC-1α),^10^ vascular endothelial growth factor (*VEGF*),^11^ and troponin T (*TNNT1*, ssTnT),^12^ respectively. While *PPARGC1A*, *VEGF*, and *TNNT1* are all differentially expressed, there is also evidence that exercise impacts the expression of different transcript variants of each of these genes,^12–15^ indicating that alternative RNA splicing (AS) is impacted by exercise.^16,17^ To date, however, only one group has investigated mRNA isoform expression using short-read RNA sequencing (SRS) in skeletal muscle from individuals with varying levels of physical activity and observed associations between activity level and mRNA isoform expression.^17,18^

RNA splicing and AS are RNA processing steps that offer additional regulatory layers in gene expression. RNA splicing is mediated by a dynamic RNA-protein complex, the spliceosome, that assembles on primary transcripts (i.e., precursor messenger RNA, pre-mRNA) to remove introns and splice exons together to form the final protein-coding transcript (i.e., mature messenger RNA, mRNA).^19^ In conjunction with the spliceosome, RNA-binding proteins (RBPs) provide tissue-/context-specific regulation of splicing.^20^ While most exons of a pre-mRNA undergo constitutive splicing to form full-length variants, many undergo AS: the choice of including or excluding certain exons and/or introns in the mRNA.^21,22^ AS is a highly conserved co-/post-transcriptional mechanism that functions to generate multiple RNA transcript variants: these transcript variants may be intended for either translation to respective protein isoforms,^23,24^ non-coding RNAs,^25,26^ or degradation via RNA decay mechanisms.^27,28^ Therefore, a single gene can form several protein isoforms depending on the demands of the cells in a tissue.^23^ For instance, a significant change in AS can orchestrate a switch in the expression of certain protein isoforms. These differential use of isoforms between conditions is often referred as isoform switching (or differential isoform expression, DIE),^13,14^ which facilitates a functional change in cellular responses to stimuli. The significance of AS is evident as nearly all mammalian genes undergo AS and generate two or more protein isoforms, which vastly expands the protein-coding capacity of the genome. AS also greatly impacts the function of cells in a tissue, as well as their responses to homeostatic perturbations.^23^

Interestingly, transcriptome-wide impact of endurance exercise on AS and isoform generation is currently unknown. Previously, Tonevitsky et al.^29^ observed that acute aerobic exercise with maximal intensity in trained skiers resulted in an increased gene expression from a set of RBPs (DEAD-box helicase 17, *DDX17*; DEAD-box helicase 46, *DDX46*; heterogeneous nuclear ribonucleoprotein R, *HNRNPR*; pre-mRNA processing factor 4 kinase, *PRPF4B*; and SR protein kinase 2, *SRPK2*) that regulate the formation of the precatalytic spliceosome. Similarly, our laboratory also observed that exercise resulted in reduced protein expression of splicing factor 3B subunit 4 (SF3B4) from young sedentary mouse gastrocnemius in response to acute treadmill running.^30^ These findings indicate that a single bout of endurance exercise is sufficient to potentially instigate differential AS (DAS) and DIE of many other genes through differential expression and regulation of RBPs. As such, we hypothesized that exercise results in isoform switching of a multitude of genes, including isoform switching of RBPs that regulate splicing. Therefore, we set out to globally profile AS events (mRNA variants/DIE and exon-level events) in skeletal muscle from a single bout of submaximal endurance exercise.

## 2 MATERIALS AND METHODS

### 2.1 Short-read RNA sequencing

The human SRS data had formerly been published by two different studies: Pattamapraponont et al.^31^ and Rubenstein et al.^32^ The raw FASTQ files were downloaded from publicly accessible NCBI Gene Expression Omnibus database (RRID: SCR_005012) (accession numbers GSE87749 and GSE151066, respectively). Briefly, Pattamapraponont et al. consisted of 10 young (19-25 years) and healthy (BMI of 23.1 ± 2.8) Danish male participants with a sedentary lifestyle (hsYS), and performed a single bout of intense (80% V O_2_max) cycling (cycle ergometer) for 15 minutes; muscle biopsies of the vastus lateralis were obtained pre-(hsYS-pre) and 4 hours post-exercise (hsYS-4pe).^31^ Rubenstein et al. studied two cohorts of older individuals (65-90 years): 10 (8 males, 2 females) an active/endurance trained group (hsAA) with a BMI of 24.2 ± 4.2; nine (7 males, 2 females) and a sedentary (hsAS) group with a BMI of 28.7 ± 3.3. The participants performed a single bout of endurance exercise (cycling at 60-70% of heart rate reserve) for 40 minutes. Muscle biopsies of the vastus lateralis were obtained pre-(hsAA-pre & hsAS-pre) and 3 hours post-exercise (hsAA-3pe & hsAS-3pe).^32^ To maintain consistency for the initial processing and statistical analyses of the SRS data, only single-end FASTQ files were used from the Pattamaprapanont et al. study; similarly, only males at pre- and 3 hours post-exercise time points with paired-end FASTQ files were used from the Rubenstein et al. to maintain comparability between the two data sets (Figure 1A).

**Figure 1.**
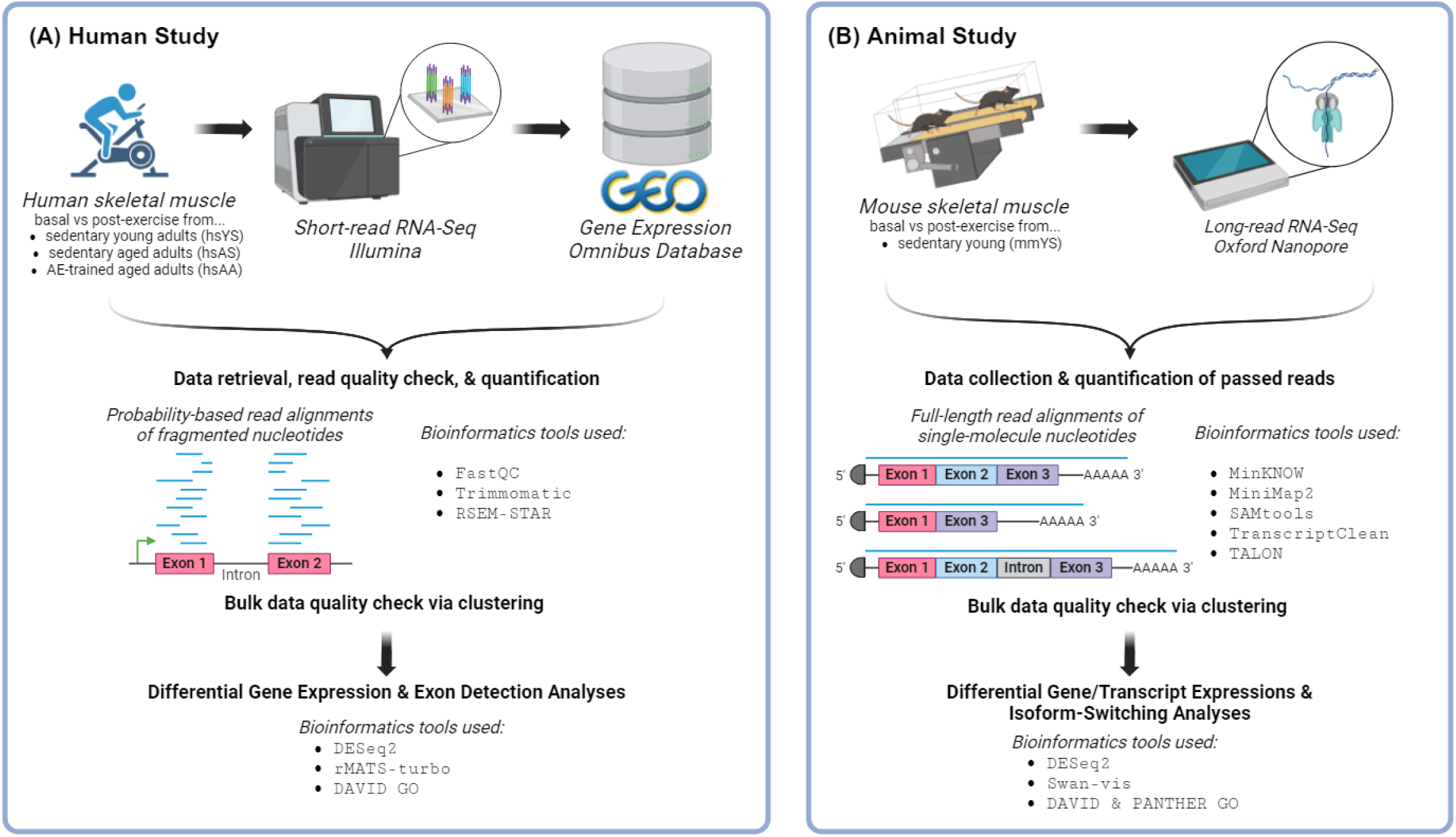
Layout of RNA-Seq study. (A) Publicly accessible human SRS data (hsYS, hsAS, and hsAA) were downloaded from NCBI Gene Expression Omnibus, preprocessed, and analyzed for differential expression. (B) Young C57BL/6 mice (mmYS) underwent an acute bout (30 minutes) of treadmill running, sacrificed, and their gastrocnemius skeletal muscle tissues were harvested for RNA sequencing cDNA library prep. Following LRS, the data were preprocessed and analyzed for differential expression. Biologically significant genes from (A) and (B) were retrieved for subsequent pathway and biased analyses.

### 2.2 Animals

All animal experiments were approved by the University of Michigan Institutional Animal Care and Use Committee (IACUC) and conducted as per the institutional guidelines. C57BL/6-Tg(TERT)C10Hode/J (approximately 8 weeks old) were purchased from The Jackson Laboratory (Bar Harbor, Maine, USA, RRID: IMSR_JAXX:006140). The mice were housed at 25°C on a 12:12-hour light-dark cycle, fed *ad libitum* laboratory mouse chow (Prolab® RMH 3000, 5P00, LabDiet by Purina, Nestlé, Vevey, Switzerland), and given free access to water. The description of the running protocol was previously published.^30^ Animals were randomly assigned to each pre-/post-exercise time points. All original animals were included without exclusion nor attrition. Investigators were not blinded to the groupings. Sample size and power analysis were determined based on our previous study.^30^ Briefly, 20 mice underwent acute treadmill running at 60% of maximal speed, approximately 75% of mouse VO_2_max according to Fernando et al.,^33^ for 30 minutes (Figure 1B,S1M). Euthanasia practices were followed as suggested by the American Veterinarian Medical Association guidelines (2020 edition): briefly, the animals were anesthetized with isoflurane, followed by cervical dislocation, and cardiac removal.

Gastrocnemius muscles were collected at pre-(mmYS-pre; 3 males, 2 females), immediately post-exercise (mmYS-ipe; 3 males, 2 females), 1 hour post-exercise (mmYS-1pe; 2 males, 3 females), and 24 hours post-exercise (mmYS-24pe; 3 males, 2 females). Gastrocnemius muscles were flash frozen in liquid nitrogen and stored at –80°C for subsequent tissue processing and RNA isolation.

### 2.3 Tissue processing, RNA isolation, and poly(A)-mRNA isolation

The gastrocnemius tissues were first powdered with a mortar and pestle in liquid nitrogen, then either processed immediately or stored at –80°C, as described in Slusher et al.^30^ Total RNA from powdered tissues were isolated using the RNeasy Plus Universal Mini Kit (Qiagen, Cat# 73404) according to manufacturer instructions, and stored at –80°C. Isolated RNA was quantified using the NanoDrop™ 2000 spectrophotometer and assessed using the Qubit™ RNA IQ Assay Kits (Invitrogen, Cat# Q33221) with the Qubit 4 Fluorometer. Subsequently, 1.2 μg of total RNA was used to isolate poly(A)-containing RNA with the NEBNext® Poly(A) mRNA Magnetic Isolation Module (New England BioLabs, Cat# E7490S) accordingly. Poly(A)-mRNA isolates were then quantified with Agilent Bioanalyzer (RRID: SCR_018043), NanoDrop™ 2000 spectrophotometry, and Qubit Fluorometry.

### 2.4 Long-read RNA sequencing

1 ng of poly(A)-mRNA isolates were used to synthesize barcoded PCR-cDNA libraries (Oxford Nanopore Technologies®, ONT; Cat# SQK-PCB109) according to the manufacturer instructions. Subsequently, the prepared PCR-cDNA libraries were quantified, multiplexed at equal molar concentrations, loaded onto a MinION™ R9.4.1 Flow Cell (ONT, Cat# FLO-MIN106) and sequenced (long-read RNA sequencing, LRS) for 72 hours using the MinION™ Mk1C sequencer (ONT, Cat# MIN-101C) (Figure 1B). During the sequencing period, the flow cell was “washed” with the Flow Cell Wash Kit (ONT, Cat# EXP-WSH004) and reloaded with fresh multiplexed libraries as needed.

### 2.5 Data preprocessing

For the SRS retrieved data, the quality of the raw FASTQ files were checked and trimmed to the minimum read length of 25 bases. Subsequently, RSEM-STAR^34,35^ was used to align and quantify the trimmed reads based on the human GENCODE release 42 (GRCh38.p13). Matrices of “counts” (estimation of expected read counts), “abundance” (TPM-normalized counts), and “length” (effective transcript lengths) were generated for statistical analyses (hsYS: Figure S1A,E,I; hsAS: Figure S1B,F,J; and hsAS: Figure S1C,G,K).

For the LRS generated data, raw FASTQ (converted/basecalled from FAST5 with Guppy MinKNOW v22.03.4; ONT, RRID: SCR_023196) files, which passed the MinKNOW quality check with minimum *q*-score threshold of 8, were used for read alignment. Minimap2 v2.14^36^ (RRID: SCR_018550) and SAMtools v1.13^37^ (RRID: SCR_002105) were used to align the passed FASTQ files into SAM format; the mouse GENCODE release M29 (GRCm39) genome sequence assembly was used for the alignment step. TranscriptClean^38^ was used to sort and “clean” the aligned SAM files. TALON^39^ was used to quantify reads and generate an abundance matrix (coverage = 0.75, identity = 0.8) in GTF format for statistical analyses (Figure S1D,H,L).

### 2.6 Differential gene expression and differential alternative splicing analyses

For the SRS-retrieved data, TXImport^40^ was used to import the multidimensional RSEM-STAR quantification matrix arrays at the gene-level to the R environment. Subsequently, DESeq2^41^ was used to perform gene-level differential expression (differential gene expression, DGE) analyses with log_2_ fold changes (log_2_FC) and respective statistics (*p*-values adjusted for FDR with default Benjamini-Hochberg method, BH; *p*-adj) on the quantification matrix arrays, and enriched for significantly up-/down-regulated genes (Figure 2A–E). IsoformSwitch-AnalyzeR^42^ was initially used (Figure S2) to identify genes with isoform-switching events with absolute difference in isoform fraction > 10% and *p*-adj < .05. Splicing type resulting in the isoform switches were annotated as skipped exon (SE), alternative 5’ splice site (A5SS), alternative 3’ splice site (A3SS), mutually exclusive exons (MXE), multiple exon-skipping (MES), retained introns (RI), alternative transcription start site (ATSS), and alternative transcription termination site (ATTS). Additionally, rMATS-turbo v4.1.2^43,44^ was used to directly analyze exon-level splicing events with difference in percent spliced in (ΔΨ or DPSI) and respective statistics on the trimmed FASTQ files, and enriched for genes with significant splicing events annotated as SE, A5SS, A3SS, MXE, and RI. The transcript-level rMATS data frames were filtered based on the absolute ΔΨ > 10% and *p*-adj < .05 for the significant splicing overlap analyses and for visualizing the proportions of splicing profile (Figure 4).

**Figure 2.**
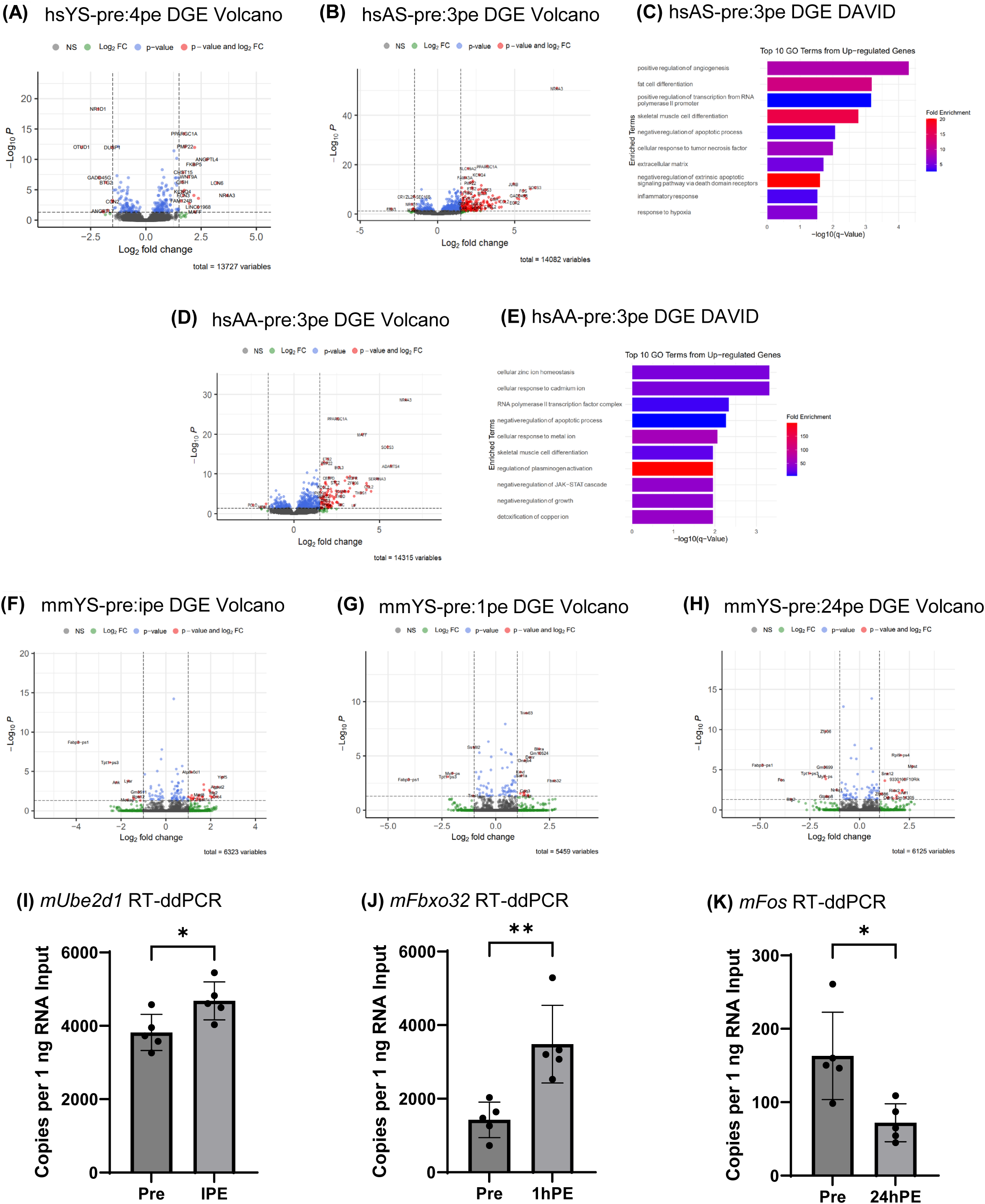
Differential gene expression (DGE) of hsYS (A), hsAS (B,C), hsAA (D,E), and mmYS (F,G,H) in the skeletal muscle comparing pre- and post-acute exercise, with LRS validated using droplet digital RT-PCR (RT-ddPCR) validations of LRS DGE data (I,J,K). (A) Volcano plot of hsYS-pre and hsYS-4pe DGE. (B) Volcano plot of hsAS-pre and hsAS-3pe DGE. (C) Top 10 GO terms from significantly up-regulated genes of hsAS-pre and hsAS-3pe DGE. (D) Volcano plot of hsAA-pre and hsAA-3pe DGE. (E) Top 10 GO terms from significantly up-regulated genes of hsAA-pre and hsAA-3pe DGE. (F) Volcano plot of mmYS-pre and mmYS-ipe DGE. (G) Volcano plot of mmYS-pre and mmYS-1pe DGE. (H) Volcano plot of mmYS-pre and mmYS-24pe DGE. (I) DGE of mmYS-pre and mmYS-ipe was validated with *mUbe2d1* using RT-ddPCR. (J) DGE of mmYS-pre and mmYS-1pe was validated with *mFbxo32* using RT-ddPCR. (K) DGE of mmYS-pre and mmYS-24pe was validated with *mFos* using RT-ddPCR. *, *p* < 0.05 compared to mmYS-pre. **, *p* < 0.01 compared to mmYS-pre. All experiments were conducted with *n* = 5 per cohort.

**Figure 3.**
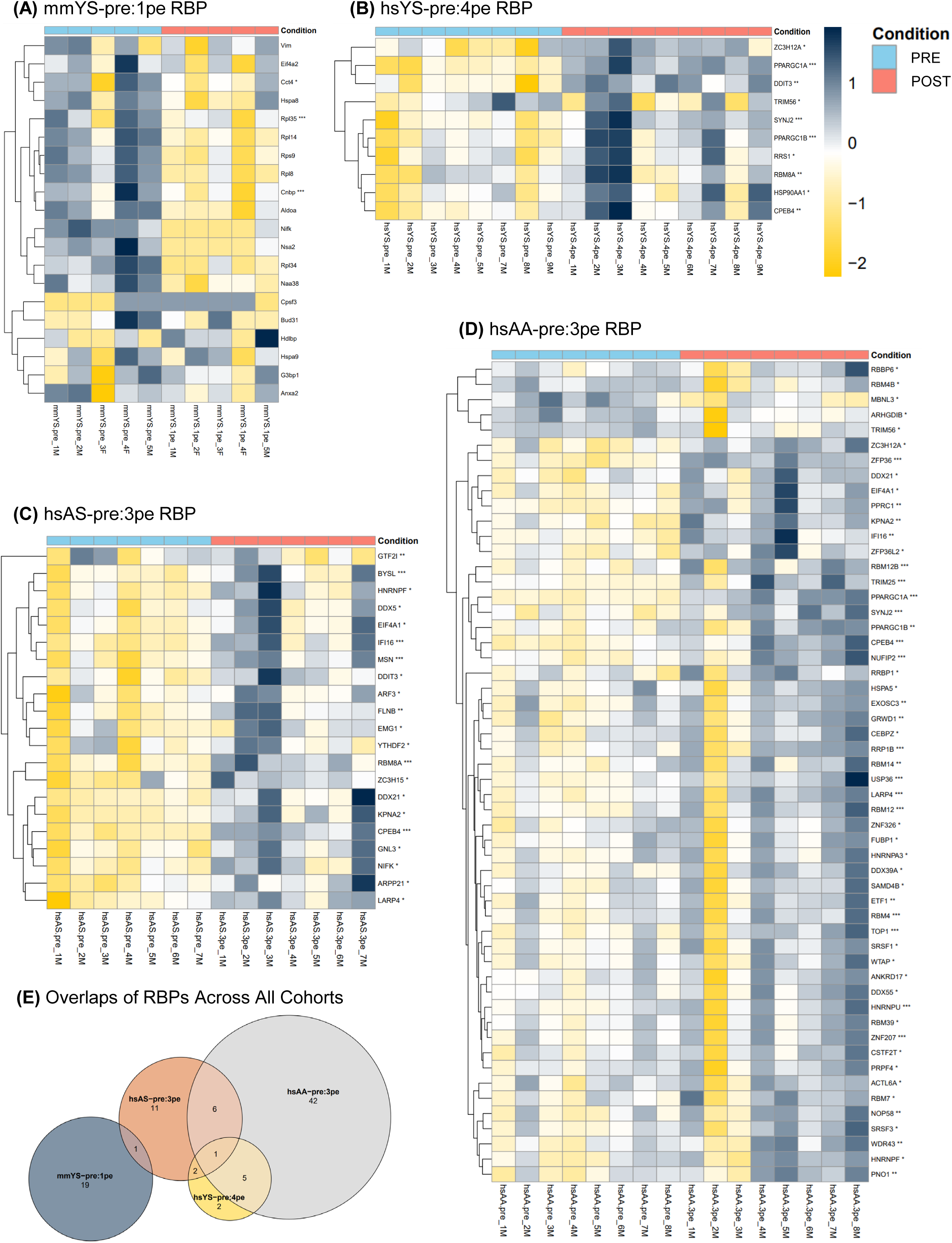
89 RBPs with significant DGE (|log_2_FC| > 0.5; raw *p* < .05 for mouse, *p*-adj < .05 for human) in response to an acute bout of exercise in skeletal muscle were visualized on the individual subject basis (raw counts converted to relative Z-scores) in mmYS (A), hsYS (B), hsAS (C), and hsAA (D), and the RBP genes were overlapped across the cohorts to find common RBPs (E).

**Figure 4.**
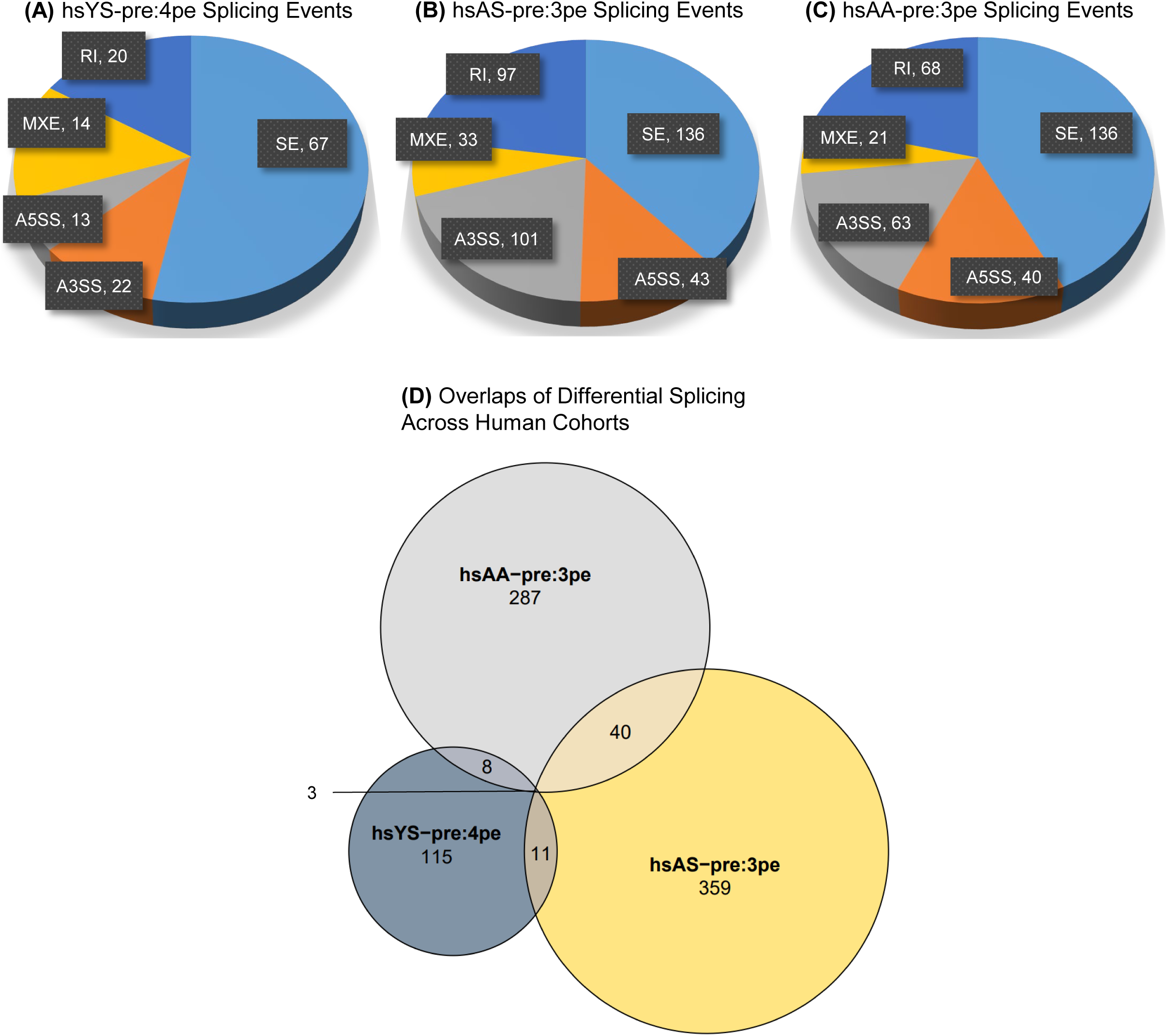
Differential splicing events in acutely exercised young and aged skeletal muscle with and without training. (A) Level of potential splicing change detected between hsYS-pre and hsYS-4pe, categorized into 5 common event types (SE, RI, MXE, A5SS, and A3SS). (B) Level of potential splicing change detected between hsAS-pre and hsAS-3pe, categorized into 5 common event types. (C) Level of potential splicing change detected between hsAA-pre and hsAA-3pe, categorized into 5 common event types. (D) Euler overlap of genes with differences in potential splicing events detected pre- and post-acute exercise across the three human SRS cohorts.

For the LRS-generated data, Swan v2.0^45^ was used to generate data frames with log_2_FC and respective significance statistics at the gene- and transcript-levels. Based on known genes that were detected in at least half of the samples, the counts matrix was generated for the downstream differential analyses. The corresponding DGE (gene-level) log_2_FC and respective statistics from Swan were used for data visualizations (Figure 2F,G,H). Similarly, DIE analyses generated absolute difference in percentage isoform use (ΔΠ or DPI) to determine isoform-switching data, whereby respective statistics from Swan were filtered based on ΔΠ > 10% and *p*-adj < .05.

Functional enrichment analysis was performed on significantly up- and down-regulated genes from the DGE analyses (|log_2_FC| > 1.5 and *p*-adj < .05) using either Database for Annotation, Visualization, and Integrated Discovery (DAVID)^46^ or PANTHER^47^ to generate top 10 gene ontology (GO) terms. Enrichment analysis was only performed on data sets with greater than 80 genes observed to be significant according to experiment specific statistical levels. With the splicing and DIE data sets from both SRS and LRS data, multiple multivariate overlap analyses across varying conditions and data sets were performed to identify any common genes with potential biologically significant splicing events. Only overlapping genes with *p*-adj < .05 from bivariate hypergeometric tests were considered significant. All code scripts used for bioinformatical data analyses are available in the Deep Blue data repository (DOI will be provided upon publication) or upon request.

An in-house-derived RNA-binding protein (RBP) list was constructed. We generated this list using two sources: the census of human RBPs^48^ and a gene cards list search for RBPs using human genes. From there we combined the two lists, generating a list of ∼1700 human genes without redundancy. Then we matched the human gene IDs to rodent gene IDs and came up with 1130 rodent genes that are conserved RBPs. This list was used to analyze RBP-specific enrichment in the data sets.

### 2.7 Gel-based and droplet digital RT-PCR

1 μg of total RNA was used to synthesize cDNA using the SuperScript™ IV reverse transcriptase (ThermoFisher Scientific, Cat# 18090010), as previously described in Slusher et al.^30^ All cDNAs were diluted 1:40 (20 µL of cDNA + 780 µL nuclease-free water; ie, 1.25 ng/µL of RNA input) before use for both droplet digital PCR (ddPCR) and gel-based PCR validations, and were stored at −20°C thereafter. All cDNA samples were assayed for 18S rRNA to ensure equal loading across the cohorts using ddPCR (Figure S5A, F: 5’-GTA ACC CGT TGA ACC CCA TT-3’, R: 5’-CCA TCC AAT CGG TAG TAG C-3’). Subsequently, several genes found significant from the LRS DGE analysis were selected for validation using RT-ddPCR (Figure 2I, *mUbe2d1*, exon 4 F: 5’-TCC TCA CTG TCC ACT TTC CG-3’, exon 5 R: 5’-TCG ATA CAG TCA AAG CGG GC-3’; Figure 2J, *mFbxo32*, exon 2 F: 5’-TCA GCA GCC TGA ACT ACG AC-3’, exon 4 R: 5’-GGC AGT CGA GAA GTC CAG TC-3’; Figure 2K, *mFos*, exon 2 F: 5’-TTT ATC CCC ACG GTG ACA GC-3’, exon 2 R: 5’-TCT ACT TTG CCC CTT CTG CC-3’). Several genes found significant from the LRS isoform-switching analysis were also selected for validation using both RT-ddPCR and gel-based RT-PCR (Figure 5C, *mSirt2*, exon 1 F: 5’-CTC TTC TTG TTT CCG CTG CC-3’, exon 3 R: 5’-CTC CCT CAG TGT CCG AGT CT-3’; Figure 6F, *mHnrnpa3*, exon 3 F: 5’-GGT GGA TGC TGC AAT GTG TG-3’, intron 3 R: 5’-ACC AAA CTG AAA CAT ATC CAA GT-3’, exon 4 R: 5’-TTT CCC ACT CTG CCT GTC TT-3’; Figure 6H, *mHnrnpa1*, exon 7 F: 5’-GGG GAT GGC TAT AAT GGA TTT GGC-3’, exon 9 R: 5’-GCA AAG TAC TGG CCT CCA CC-3’; Figure 6I, *mTia1*, exon 4 F: 5’-GCA ACA ACC CCT AGC AGT CA-3’, exon 6 R: 5’-CGC TGC TTT GAT GTC TTC GG-3’).

**Figure 5.**
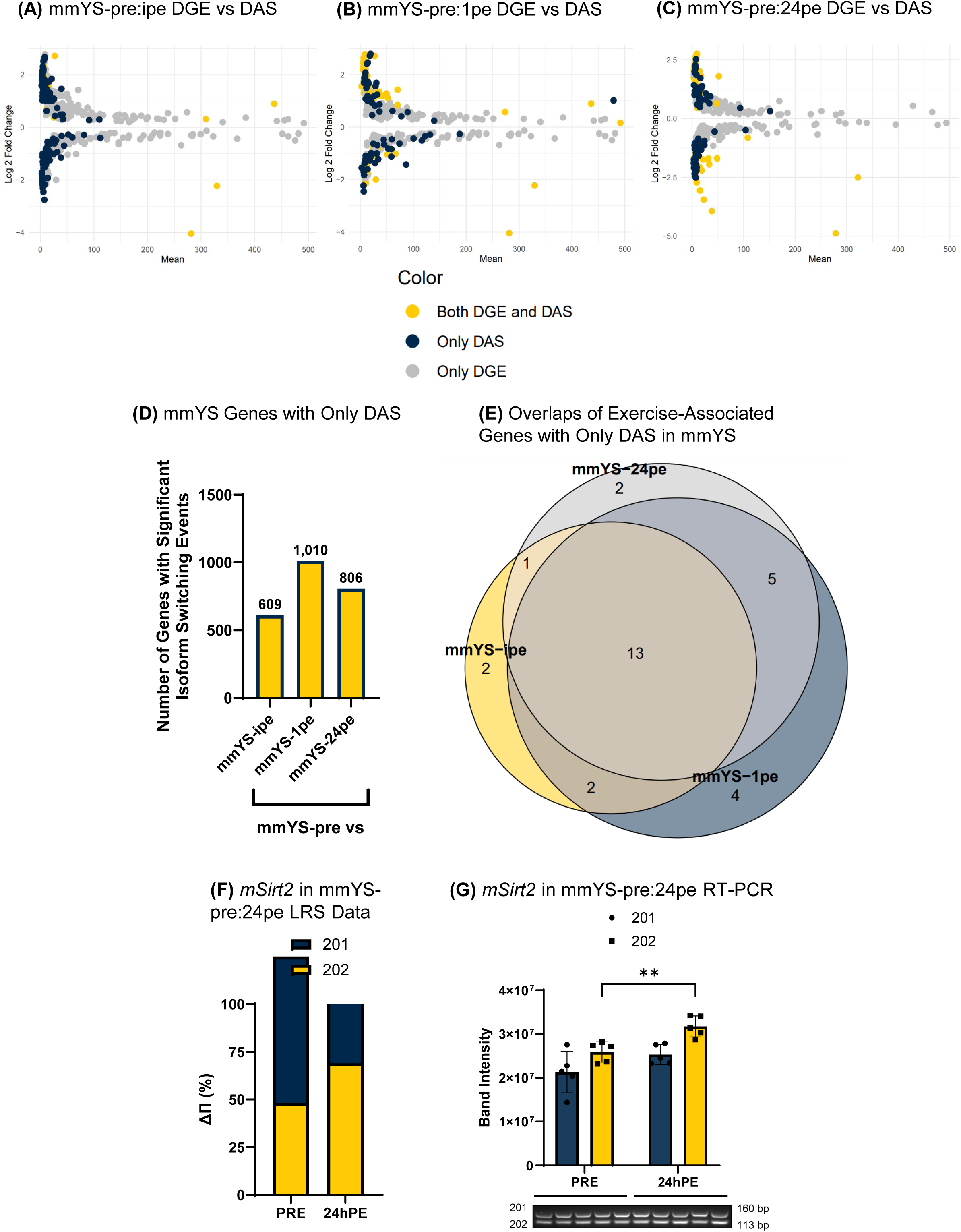
Splicing and isoform switching in acutely exercised young sedentary skeletal muscle in a temporal manner. (A) Expressional changes in genes with significant (raw *p* < .05) DGE and DAS between mmYS-pre and mmYS-ipe. (B) Expressional changes in genes with significant DGE and DAS between mmYS-pre and mmYS-1pe. (C) Expressional changes in genes with significant DGE and DAS between mmYS-pre and mmYS-24pe. (D) Isoform-switching genes with statistical significance (ΔΠ > 10%, *p*-adj < .05) but without significant DGE between pre- and post-acute exercise time points. (E) Euler overlap of exercise-associated genes with statistical significant isoform switching but without significant DGE between pre- and post-acute exercise time points. (F) DIE of *mSirt2*, an exercise-associated gene, between its 201 and 202 transcript variants between pre- and 24 hours post-exercise according to LRS data. (G) DIE of *mSirt2*-202 transcript variant between pre and 24pe was validated using gel-based RT-PCR. **, *p* < 0.01 compared to mmYS-pre. All experiments were conducted with *n* = 5 per cohort.

**Figure 6.**
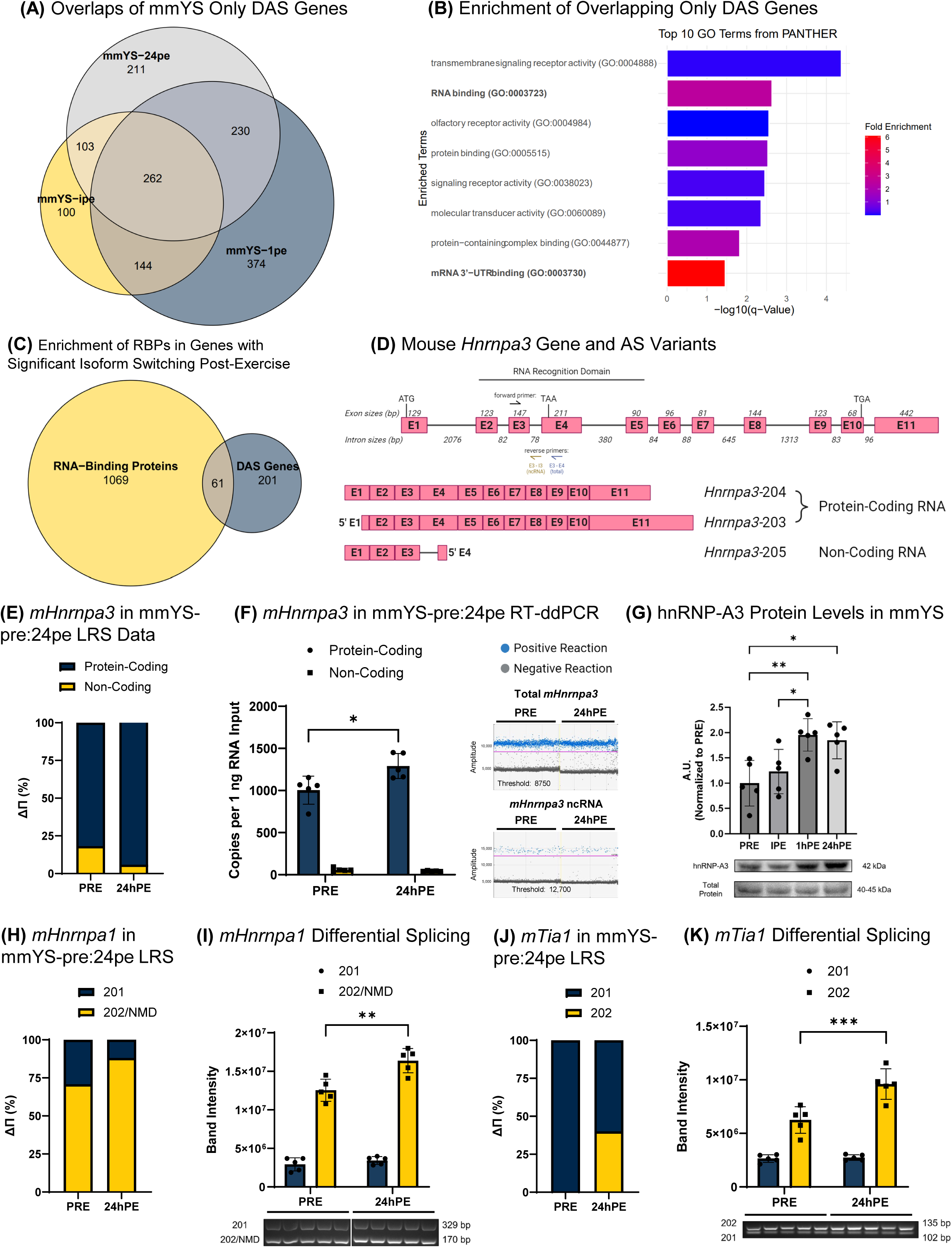
Splicing and isoform switching of RBPs in acutely exercised young sedentary skeletal muscle in a temporal manner. (A) Euler overlap of all genes with statistically significant isoform switching but without significant DGE between pre- and post-acute exercise time points found in Figure 5D. (B) Top 10 GO terms from the 262 common genes across the three post-acute exercise time points found in Figure 6A using PANTHER. (C) Overlap of the 262 common genes with in-house list of 1130 RBP genes, highlighting 61 RBP genes without significant DGE. (D) Overview of mouse *Hnrnpa3* gene loci and location of primers used for RT-ddPCR, and the exon/intron structures of transcript variants 203, 204, and 205. (E) DIE of *mHnrnpa3*, an RBP gene, between its protein-coding (all reads not assigned to 205) and non-coding (205, retained intron) transcript variants between pre- and 24 hours post-exercise according to LRS data. (F) DIE of *mHnrnpa3* protein-coding transcripts between pre and 24pe was validated using RT-ddPCR. (G) Western blot validation of significant isoform switching of hnRNP-A3 at the protein level, normalized to total protein at 40-45 kDa band detection then normalized to mmYS-pre for relative quantification in arbitrary units (AU), with representative image of western blots. (H) DIE of *mHnrnpa1*, an RBP gene, between its 201 and 202/nonsense-mediated decay transcript variants between pre- and 24 hours post-exercise according to LRS data. (I) DIE of *mHnrnpa1*-202/nonsense-mediated decay transcript variants between pre and 24pe was validated using gel-based RT-PCR. Full RT-PCR gel shown in S5B. (H) DIE of *mTia1*, an RBP gene, between its 201 and 202 transcript variants between pre- and 24 hours post-exercise according to LRS data. (K) DIE of *mTia1*-202 transcript variant between pre and 24pe was validated using gel-based RT-PCR. *, *p*-adj < 0.05 compared to mmYS-pre. **, *p*-adj < 0.01 compared to mmYS-pre. ***, *p*-adj < 0.001 compared to mmYS-pre. All experiments were conducted with *n* = 5 per cohort.

### 2.7 Protein lysate preparation, antibody validation, and western blot analyses

To examine protein expression level of one of the prominent RBPs, heterogeneous nuclear ribonucleoprotein A3 (hnRNP-A3), HEK 293 fetal kidney cells (ATCC, Cat# CLR-1573, RRID: CVCL_0045) were used as positive controls to verify cross-species specificity of the hnRNP-A3 antibody for the correct molecular weight at 42 kDa. HEK 293 blots were normalized with GAPDH (conditions listed below). Additionally, C2C12 murine myoblasts (ATCC, Cat# CRL-1772, RRID: CVCL_0188) were transfected with 30 nM of either scrambled (siCTL) or mouse *Hnrnpa3*-targeting (exon 10) siRNAs (ThermoFisher Scientific, Cat# 4390771) in Lipofectamine™ RNAiMAX transfection reagent (Invitrogen, Cat# 13778075) for 24 and 48 hours to knock-down hnRNP-A3 and thus validate the protein specificity of hnRNP-A3 antibody. All cell lines used were authenticated by the manufacturer (ATCC) and visually verified in-house. The cell pellets were resuspended in an SDS-based lysis buffer, sonicated, and centrifuged to extract the total protein lysate. Total protein lysates were prepared from gastrocnemius, as described in Slusher et al.,^30^ were used to quantify protein expression level of hnRNP-A3 in the skeletal muscle tissues. Protein concentrations were quantified using the Pierce 660 nm Protein Assay Kit (ThermoFisher Scientific, Cat# 1861426). These protein lysates from cell lines and the animal-derived tissues were then prepared using 2X Laemmli loading buffer (Bio-Rad, Cat#1610737) and boiled for 10 minutes at 95°C.

All protein lysates from cells and tissues were resolved by SDS-polyacrylamide gel electrophoresis at 16 mAmp per gel for 1 hour, transferred to Mini 0.2 μm polyvinylidene fluoride membranes using the Trans-Blot® Turbo™ Transfer System (Bio-Rad, Cat# 1704150) with the rapid 7-minute mixed molecular weight transfer protocol. To verify cross-species specificity of the hnRNP-A3 antibody, protein lysates of wild-type HEK 293 was used. To quantify bands from the wild-type HEK 293 samples, specifically, loading control was normalized with GAPDH (Cell Signaling Technology, Cat# 2118, RRID: AB_561053; 1:1000 dilution in 5% bovine serum albumin with 1X PBST) (Figure S6A). To validate protein specificity of the hnRNP-A3 antibody, protein lysates of C2C12 myoblasts treated with either siCTL or *Hnrnpa3* siRNA were used. To quantify bands from the C2C12 myoblasts, specifically, loading control was normalized with total protein quantification of bands at 40-45 kDa using Pierce™ Reversible Protein Stain Kit (ThermoFisher Scientific, Cat# 24585) (Figure S6B,C).

HnRNP-A3 from these cell lysates was detected with a rabbit polyclonal antibody (Abcam, Cat# ab78300, RRID: AB_2041662; 1:1000 dilution factor in 5% bovine serum albumin with 1X PBST and 0.02% sodium azide). To quantify bands from the skeletal muscle tissue, specifically, loading control of protein lysates from the skeletal muscle tissue was normalized with total protein quantification of bands at 40-45 kDa using Ponceau S stain (Sigma-Aldrich, Cat# P3504) and detected for hnRNP-A3 with the validated polyclonal antibody (Figure 6G,S6D). All blots were imaged with Bio-Rad ChemiDoc XRS+ Molecular Imager (RRID: SCR_019690) and quantified with Bio-Rad Image Lab software. Values from either the reversible Pierce-/Ponceau S-stained blots (tissue-based lysates) or the loading control blots (cell-based lysates) were averaged and used to normalize the target protein quantification. All images that were used to quantify hnRNP-A3 expression levels are included in Figure S6D.

### 2.8 Statistical analysis

Unless otherwise noted, the unpaired Student’s *t*-test was used for two group comparisons using a statistical package (GraphPad Prism 10, RRID: SCR_002798). One-way ANOVAs were used to determine significance for multiple group comparisons of the western blot data, whereby the groups (time points) had 1 level each. Subsequently, *post hoc* adjustments were conducted with Tukey’s HSD. Statistical significance was defined as a *p* < .05.

## 3 RESULTS

### 3.1 RNA-Seq identified differential gene expression in acutely exercised skeletal muscle with and without training

We analyzed gene expression from public SRS data and compared these findings to our acute exercise LRS study. DGE of vastus lateralis in hsYS-pre and hsYS-4pe was analyzed with DESeq2 (Figure 2A): out of 13727 total detected genes, 19 were significantly up-regulated (log_2_FC > 1.5, *p*-adj < .05) including *PPARGC1A* (i.e., PGC-1α), nuclear receptor subfamily 4 group A member 3 (*NR4A3*), and heat shock protein family A member 1A/B (*HSPA1A/B*, HSP70/72), while 10 were significantly down-regulated (log_2_FC < −1.5, *p*-adj < .05). There were insufficient numbers of DGE genes, therefore it was not appropriate to conduct functional enrichment analyses. DGE of vastus lateralis in hsAS-pre and hsAS-3pe was analyzed (Figure 2B): out of 14082 total detected genes, 201 were significantly up-regulated (log_2_FC > 1.5, *p*-adj < .05) including *PPARGC1A*, *NR4A3*, and *HSPA1B*, while 9 were significantly down-regulated (log_2_FC < −1.5, *p*-adj < .05). We took all significantly up-regulated genes and performed functional enrichment analysis (DAVID, statistical over-enrichment test). The hsAS-pre:3pe revealed 204 terms (*p*-adj < .05; Figure 2C). Similarly, DGE of vastus lateralis in hsAA-pre and hsAA-3pe was analyzed (Figure 2D): out of 14315 total detected genes, 112 were significantly up-regulated (log_2_FC > 1.5, *p*-adj < .05) including *PPARGC1A*, *NR4A3*, nuclear receptor subfamily 4 group A member 1 (*NR4A1*), and *HSPA1B*, while 4 were significantly down-regulated (log_2_FC < −1.5, *p*-adj < .05). We took all significantly up-regulated genes and performed functional enrichment analysis (DAVID, statistical over-enrichment test). The hsAA-pre:3pe revealed 149 terms (*p*-adj < .05; Figure 2E). GO terms that were common between the hsAS and hsAA groups were skeletal muscle cell differentiation (GO:0035914) and negative regulation of apoptotic process (GO:0043066). The major insights we are able to draw from these data are that there is a different response to acute aerobic exercise based on training status: DGE analyses illustrated different up-regulated genes post-exercise in the older active individuals with few GO terms in common between the two older cohorts.

DGE of gastrocnemius in mmYS-pre and mmYS-ipe, mmYS-pre and mmYS-1pe, and mmYS-pre and mmYS-24pe were analyzed (|log_2_FC| > 1.5, *p*-adj < .05). At ipe, 6323 total genes were detected, and we observed 23 differentially expressed genes (6 down-regulated and 17 up-regulated); at 1pe, 5459 total genes were detected, and we observed 13 differentially expressed genes (4 down-regulated and 9 up-regulated); and at 24pe, 6125 total genes were detected, and we observed 31 differentially expressed genes (16 down-regulated and 15 up-regulated). Since this was a conservative statistical cutoff, we relaxed the parameters to include more genes in our analysis (|log_2_FC| > 1.0, *p*-adj < .05; Figure 2F, G and H). With these new statistical parameters, at ipe, we detected 46 genes that reached statistical significance (13 down-regulated and 33 up-regulated); at 1pe, we detected 32 genes that reached statistical significance (8 down-regulated, 24 up-regulated); and at 24pe, we detected 49 genes that reached statistical significance (26 down-regulated, 23 up-regulated). To validate our data sets we selected one gene at each time point for RT-PCR validation. Ubiquitin-conjugating enzyme E2 D1 (*mUbe2d1*; LRS data: log_2_FC = 0.75, *p*-adj = 0.0002), F-box protein 32 (i.e., atrogin-1, *mFbxo32*; LRS data: log_2_FC = 2.7, *p*-adj = 0.002), and fos proto-oncogene (*mFos*; LRS data: log_2_FC = –3.9, *p*-adj = 0.0002) at ipe, 1pe, and 24pe, respectively. Using RT-ddPCR, we validated all three selected genes at each timepoint (*mUbe2d1* was significantly up-regulated at ipe [22.6% increase, *t*-test *p* = 0.03); *mFbxo32* was up-regulated at 1pe [144% increase, *t-*test *p* = 0.004); and *mFos* was down-regulated at 24pe [55.9% decrease, *t-*test *p* = 0.01]; Figure 2I,J,K, respectively). Genes commonly observed to be differentially expressed in acute endurance exercise studies such as *mHspa8* (i.e., HSP70)^49^ were observed in this analysis. Interestingly, we observed that exercise impacted on genes related to protein degradation, such as *mFbxo32* and *mUbe2d1*, indicating that exercise seemed to enhance clearing damaged proteins.^50,51^

We did not observe the common acute exercise-induced genes such as NRFs or PGC-1α in our LRS data like we did using the publicly available human SRS data. A potential reason for this is the low expression of these transcription factors, as well as the timing of muscle sampling post-exercise in our mouse study (1 hour versus 3-4 hours post exercise in the humans).

### 3.2 Differentially expressed RNA-binding proteins in acutely exercised skeletal muscle with and without training

Using a list of 1130 RBP genes generated in house (see methods above for details on how this list was curated), select RBP genes were included in the heatmap if it was significantly differentially expressed within a cohort. Significant up- and down-regulated RBPs (log_2_FC > 0.5, log_2_FC < −0.5, raw *p* < .05) in response to an acute bout of exercise from all four cohorts (mmYS, hsYS, hsAS, and hsAA) were queried, identifying 89 genes across all four groups (Figure 3A–D, *p*-adj < .05 indicated with asterisks). The mmYS cohort had 3 RBPs that were significantly down-regulated with *p*-adj < .05 in response to acute exercise (Figure 3A). We show an additional 17 RBP genes (10 up-regulated, 7 down-regulated, raw *p* < .05) with less conservative statistical parameters (Figure 3A). The hsYS cohort had a total of 9 and 1 RBPs were significantly up- and down-regulated, respectively, in response to acute exercise (Figure 3B). The hsAS cohort had a total of 20 and 1 RBPs were significantly up- and down-regulated, respectively, in response to acute exercise (Figure 3C). The hsAA cohort had a total of 50 and 4 RBPs were significantly up- and down-regulated, respectively, in response to acute exercise (Figure 3D). Finally, we overlapped the RBPs to determine conserved and age-related changes in RBP expression between cohorts (Figure 3E). We did not observe any overlapping genes across the mice and human cohorts. This could indicate that there is not a well-conserved response to endurance exercise and changes in the mRNA expression of RBPs or this could indicate differences in the exercise protocols and muscles sampled between the cohorts (gastrocnemius for the mice versus quadriceps for the humans). There were 6 overlapping RBPs that were significant between the aged cohorts using hypergeometric testing (6.0-fold over-enrichment, *p* = 0.0002). There were also 5 overlapping RBPs that were significant between the YS and AA cohorts using hypergeometric testing (10.5-fold over-enrichment, *p* = 4.4*e*^-5^). There were also 2 overlapping RBPs that were significant between hsYS and hsAS cohorts using hypergeometric testing (11.3-fold over-enrichment, *p* = 0.02).

Cytoplasmic polyadenylation element binding protein 4 (CPEB4) was a common up-regulated RBP gene among the human cohorts. CPEB4 is a protein mainly found in the cytoplasm that is a sequence-specific RBP and binds to uridine-rich sequence elements (consensus sequence 5’-UUU UUA U-3’) in the 3’-untranslated region of mRNAs in the cytoplasm regulating their stability/degradation and their translatability. Specifically, during energetic stress this protein blocks global translation while preserving translation of essential proteins.^52^ CPEB4 is involved in muscle regeneration and metabolism and in response to exercise and caloric restriction.^53,54^ Overall, these data indicate that acute endurance exercise results in a mild response in terms of gene expression changes of RBPs in young organisms; however, that the response of gene expression changes of RBPs is much more robust in older individuals.

### 3.3 SRS identified differential splicing events in acutely exercised young and aged skeletal muscle with and without training

Isoform-switching and DIE analyses in human skeletal muscles following acute exercise were first attempted with IsoformSwitchAnalyzeR using only the SRS data. Firstly, DIE revealed the most common AS event type of each human cohort (Figure S2A,B,C). Isoform switching was also determined: compared to the respective baselines, 10 genes had significant isoform-switching events in hsYS, 15 in hsAS, and 21 in hsAA from an acute bout of exercise (Figure S2D,E). These minimal changes were attributed to the read counts and paired-end versus non-paired end reads of the SRS data sets. Subsequently we choose to utilize an exon splicing data analysis pipeline to glean additional alternative splicing information from the SRS data.

#### Splicing events at the exon level were analyzed with rMATS, using only the SRS data

The most common event across the three human cohorts was SE followed by RI. The hsYS cohort revealed 67 SE events (Figure 4A); the hsAS revealed 136 SE events (Figure 4B); the hsAA revealed 136 SE events (Figure 4C); all with statistical significance (*p*-adj < .05). When comparing the ratio of splicing event types between the younger humans and the older humans, it is interesting to note that exercise induced more A5SS and A3SS in older individuals compared to younger, though this could potentially be due to the differences in the exercise protocols between the studies as well. Also, these data illustrate a considerable difference in number of splicing events in skeletal muscle between the younger and aged cohorts (younger cohort had 127 total events compared to greater than 320 events in both aged cohorts). Overall, exercise impacts differential exon usage, especially SE events, whereby age and exercise appear to interact to modify differential exon usage to a greater extent.

To determine if there were specific pathways that were enriched by post-exercise splicing events as determined by rMATS, we took all the genes with statistically significant differential splicing events (ΔΨ > 10%, *p*-adj < .05) and performed functional enrichment analyses (DAVID and PANTHER, statistical over-enrichment test, GO molecular function complete, Fisher’s exact test). 115 genes from the hsYS-pre:4pe revealed 1 term, protein binding GO:0005515. The same test was repeated for hsAS-pre:3pe (52 enriched terms, *p*-adj < .05; Figure S3A) and for hsAA-pre:3pe (30 enriched terms, *p*-adj < .05; Figure S3B). Protein binding was a term that was in common across all three groups. GO terms that were common between the hsAS and hsAA groups were catalytic activity (GO:00033824), binding (GO:0005488), and molecular adaptor activity (GO:0060090). The major insights we are able to draw from these data are that there is a different response based on age and training status, which is also displayed above in Figure 3 exploring which RBPs are differentially expressed. These data are in alignment with previous research.^16,55^

The genes with statistically significant differential splicing events were also compared to observe any overlaps (Figure 4D). Out of 115, 359, and 287 known genes from the hsYS, hsAS, and hsAA cohorts, respectively (unique gene IDs; some genes had multiple splicing events, but each gene ID was only entered once in the overlap analysis), 11 genes overlapped between hsYS and hsAS, 8 genes between hsYS and hsAA, 40 genes between hsAS and hsAA, and 3 genes across all. A bivariate hypergeometric test between hsAA and hsAS cohorts revealed significant over-enrichment compared to expectations (40 genes in common, 1.984-fold over-enrichment, *p* = 2.62*e*^-5^), the other overlapping genes were not statistically significant beyond random chance from the hypergeometric testing. The three genes with overlapping splicing events between the human cohorts were ras homolog family member T2 (*RHOT2*), mitochondrial arginyl-tRNA synthetase 2 (*RARS2*), and AMP-activated protein kinase non-catalytic subunit gamma-2 (*PRKAG2*, γ2-AMPK).

RHOT2 is an outer mitochondrial membrane GTPase that plays a role in mitochondrial trafficking and fusion-fission dynamics, with little information on the impact of exercise on this gene or protein product. RARS2 is a mitochondrial tRNA synthetase that plays an important role in the translation of mitochondrially encoded proteins. There is little research about the role of this gene related to exercise or skeletal muscle, but mutations in this gene are associated with neurological disease and lactic acidosis.^56,57^ The final overlapping gene, *PRKAG2*, is a part of a well-known enzyme, AMPK, associated with exercise responses. *PRKAG2* encodes for the non-catalytic γ2 subunit with 35 transcript variants that can form 12 different protein isoforms, 7 transcripts that are degraded by nonsense-mediated decay, and the remaining either lack on open reading frame or contain a retained intron that lacks a protein product. This subunit is important in binding AMP and regulating activity levels of the catalytic subunits.^58^ Further mutations in this gene cause heart issues and metabolic issues in humans.^59,60^ In terms of the functional role of the AS variants that give rise to protein isoforms of γ2-AMPK, little information is published in the literature. This represents an area ripe for future investigation in the field of exercise physiology and metabolic health. When we searched to see if the event was the same splicing event across the cohorts, we found that it was an MXE event in the hsYS cohort, while A3SS in the older hsAS and hsAA cohorts. This indicates that exercise impacts the splicing of this gene, but the events shift from an MXE event to an A3SS either due to aging or due to differences in the exercise protocols. This will have to be determined in future studies. While the γ2-AMPK is not the major gamma subunit studied or expressed in skeletal muscle, its complex regulation raises the possibility that this minor subunit plays important functional roles, and warrants future investigation.

We next wanted to determine how many RBPs had significant exon splicing events post exercise in the human cohorts. Using our RBPs list (described above) we observed 12 RBPs, 32 RBPs, and 20 RBPs had significant splicing events in hsYS, hsAS, and hsAA, respectively. Notable RBPs with significant splicing events in the hsYS cohort include heterogeneous nuclear ribonucleoprotein A2/B1 (hnRNP-A2/B1),^61^ RNA-binding Fox-1 homolog 2 (RBFOX2),^62^

RNA-binding motif protein 3 (RBM3),^63^ and cold-inducible RNA-binding protein (CIRBP),^64^ all of which have been related to muscle phenotypes in the literature. In the hsAS cohort, some interesting RBPs with splicing events include several ATP-dependent RNA helicases (DDX17, DDX23, and DDX5), components of the RNA exosome (EXOSC2, EXOSC3, EXOSC8), and SRPK2, among others. In the hsAA cohort, CLK4, RBM39, DDX17, and SRPK2 are commonly studied RBPs with significant splicing events post-exercise. To determine if there were any common RBPs across the cohorts, we overlapped the significant RBPs. This analysis revealed no overlapping genes between all three cohorts, and the only pairwise comparison with any overlapping genes was hsAA with hsAS, which revealed *SRPK2* and *DDX17*. These genes are interesting as they are both ATP-dependent enzymes with the potential to trigger downstream impacts on splicing regulation and they may be impacted by acute endurance exercise.

### 3.4 LRS identified differential gene expression, differential alternative splicing, and isoform switching in young mice during recovery from acute exercise

Significant DGE and DAS (SE and RI only) of gastrocnemius in mmYS were analyzed using Swan (log_2_FC ≠ 0, ΔΠ > 10%, raw *p* < .05) to contrast how expressional changes of transcription and AS differed across the three time points in recovery from acute exercise. Based on mean normalized counts, mmYS-ipe compared to mmYS-pre illustrated DGE only (313, 63.9%), DAS only (159, 32.4%), both DGE and DAS (18, 3.7%) profiles (Figure 5A); mmYS-1pe compared to mmYS-pre illustrated DGE only (254, 68.6%) DAS only (39, 10.5%) both DGE and DAS (77, 20.8%) profiles (Figure 5B); and mmYS-24pe compared to mmYS-pre illustrated DGE only (254, 65.5%), DAS only (50, 12.9%), both DGE and DAS (84, 21.6%) profiles (Figure 5C). These data indicate that exercise not only perturbs transcript expression but also impacts DAS in combination with transcriptional changes and independent of transcriptional changes. This reveals a new source of exercise-induced responses and adaptations that need to be explored for their functional relevance in the physiology of skeletal muscle and in their roles in exercise-induced health benefits.

Isoform switching (i.e., differential isoform expression, DIE) in gastrocnemius of mmYS-ipe, mmYS-1pe, and mmYS-24pe compared to mmYS-pre was analyzed with Swan. A total of 609, 1010, and 806 transcripts had significant isoform switching events (ΔΠ > 10%, *p*-adj < .05) at each timepoint, respectively (Figure 5D). We investigated our isoform switching data for known exercise associated genes. We generated a list of 60 well-established exercise-associated genes and overlapped with the list of genes that had significant isoform-switching events at each time point. We observed isoform switching in 18, 24, and 21 of those 60 genes in ipe, 1pe, and 24pe, respectively. According to hypergeometric testing each of these overlapping gene sets is highly over enriched compared to random expectations (ipe, 20.3-fold over-enrichment, *p* = 5.2*e*^-19^; 1pe, 17.6-fold over-enrichment, *p* = 4.6*e*^-24^; and 24pe, 17.4-fold over-enrichment, *p* = 6.5*e*^-21^). Genes including sirtuin 2 (*mSirt2*), myosin heavy chain IIb (*mMyh4,* MHC-IIb), *mNr4a1*, and myocyte-specific enhancer factor 2C (*mMef2c*) were observed to have significant DIE across all three time points post exercise (there were 13 genes in common across all three groups).

We used RT-PCR to validate the splicing event observed in *mSirt2.* The LRS data showed that there was an exon skipping event of exon 2 of Ensembl transcript *mSirt2*-201 resulting in a 22% increase in *mSirt2*-202 at 24pe (Figure 5F). We designed primers for exons 1 and 3 of *mSirt2*-201 and we were able to quantify exon skipping. When exon 2 of *mSirt2*-201 is skipped this produces *mSirt2*-202. We were able to show that there were no changes in transcript levels of *mSirt2*-201 at 24pe compared to pre (14.6%, *p* = n.s.); however, there was a significant increase in *mSirt2*-202 at the same time points (24.5% *p* = 0.005) validating the isoform-switching event post-exercise (Figure 5G). We queried if *SIRT2* had a significant exon-level splicing event post-exercise in the rMATS data of the human cohorts, but did not observe any significant events. This could be due to the different timing post-exercise sample collection (3-4 hours for the humans compared to 24 hours for the mice). There is limited data on the functional differences between *mSirt2* transcripts 201 and 202. Future studies would be required to determine the function and regulation of the exon 2 skipping event. Overall, our *mSirt2* data indicate that genes commonly associated with exercise responses and adaptations undergo isoform-switching events that may be important to investigate their physiological relevance in future mechanistic follow-up studies.

Furthermore, 144 (10.1%) overlapped between mmYS-ipe and mmYS-1pe, 103 (7.2%) between mmYS-ipe and mmYS-24pe, 230 (16.2%) between mmYS-1pe and mmYS-24pe, and 262 (18.4%) across all three groups (Figure 6A). When unique gene identifiers of significant isoform-switching genes at each time point (as described above) were queried for enriched pathways using PANTHER, this revealed several common GO terms including RNA binding (top 10 GO terms are shown in Figure S4A,B,C). When the 262 common genes across time points post-exercise were independently queried for enriched pathways, revealing enrichment of GO terms including RNA binding (GO:0003723), mRNA 3’-untranslated region (UTR) binding (GO:0003730), transmembrane signaling receptor activity (GO:0004888), and molecular transducer activity (GO:0060089) (top 10 GO terms are shown in Figure 6B). Within these 262 common genes, 61 were RBPs as identified by overlapping the 262 gene identifiers with a list of 1130 RBP genes generated in house (Figure 6C) and hypergeometric testing revealed that this was significantly over-enriched compared to expectations (24.4-fold over-enrichment, *p* = 5.1*e*^-69^).

Several top candidate RBP genes, heterogeneous nuclear ribonucleoprotein A3 (*mHnrnpa3*), A1 (*mHnrnpa1*), and T-cell intracellular antigen 1 (*mTia1*) were chosen for validation due to the abundance of each gene, and ability to design RT-PCR primers to detect the splicing event. The LRS data showed that there was a significant isoform-switching event of ΔΠ = 10% increase in the protein-coding transcripts at 24pe (Figure 6E). We designed primers for exons 3 and 4 of *mHnrnpa3* (to detect transcripts 203 and 204) and intron 3 to quantify intron retention of *mHnrnpa3*-205 (non-coding RNA) (Figure 6D). RT-ddPCR showed that there was no change in transcript levels of *mHnrnpa3*-205 at 24pe compared to pre (*t*-test *p* = n.s.); however, there was a significant increase in protein-coding *mHnrnpa3* at the same time points (28.5%, *t*-test *p* = 0.02) validating the isoform-switching event post-exercise (Figure 6F).

Western blots showed that there were significant increases of hnRNP-A3 at the protein level post-exercise (pre:ipe, 23.1%, *p*-adj = n.s.; pre:1pe, 95.6%, *p*-adj = 0.008; pre: 24pe, 84.9%, *p*-adj = 0.02; Figure 6G). The LRS data also showed that there was a significant DIE event in *mHnrnpa1*, resulting in an overall 24% increase in *mHnrnpa1*-202 (SE) and *mHnrnpa1*-203 (nonsense-mediated decay) transcripts at 24pe (Figure 6H). RT-PCR showed significant increases in *mHnrnpa1*-202 and *mHnrnpa1* nonsense-mediated decay products at 24pe (30.6%, *t*-test *p* = 0.004; Figure 6I). The LRS detected a significant DIE event in *mTia1* as well, resulting in an overall 40% increase in *mTia1*-202 (SE) transcripts at 24pe (Figure 6J), validated by RT-PCR of the 202 transcripts at 24pe (53.6%, *t*-test *p* = 0.004; Figure 6K).

## 4 DISCUSSION

The present study supports that, in addition to DGE in recovery from endurance exercise, there is a robust AS response across the transcriptome in skeletal muscles from both humans and mice. Several insights from our data include the following: 1) post-exercise recovery of skeletal muscle impacts AS; 2) the pathways enriched in splicing events appear to be related to splicing regulation (i.e., RBPs) among several other pathways; 3) common exercise-associated genes are impacted by AS post-exercise, such as *PRKAG2* (γ2-AMPK) and *SIRT2*; 4) several of the common RBPs observed to have either DGE or DAS post-exercise are related to 3’-untranslated region binding (CPEB4, hnRNP-A1, and TIA1) and transcript stability; 5) there appears to be a difference in how AS is regulated by aging with older individuals having high RBP expression and more splicing changes; and 6) our mouse LRS data revealed that more splicing events may be significant 24 hours post-exercise, compared to transcriptional events being highly enriched at earlier timepoints.

Our data indicate that while some RBPs and spliceosome proteins undergo DGE in response to an acute bout of endurance exercise in untrained skeletal muscle, far more RBPs and spliceosome proteins undergo DAS and isoform switching in response to exercise. Our rMATS analysis of SRS data revealed that young individuals have splicing events in response to endurance exercise, but both sedentary and trained older individuals have a greater number of significant events compared to younger individuals. While SRS is excellent for detecting DGE and exon splicing events, it lacks the resolution to accurately detect transcript variants and/or isoform-switching events due to multi-read alignment and mapping ambiguity.^65^ Finally, our data clearly points out the strengths of LRS for the detection of full-length transcript variants and opens an entirely new area of adaptive potential in skeletal muscle to explore in response to endurance exercise.

Recent literature has indicated changes in the expression of transcript variants with aging in skeletal muscle.^16,55,66–69^ Since endurance exercise is a potent anti-aging intervention (i.e., by maintaining V O_2_max), we surmised that the response of skeletal muscle gene expression at the level of AS would differ by age and training status. Our data clearly indicate that this is likely the case (Figure 3,4). First, since previous research has shown significant RNA and protein level differences in RBPs with age, we explored a biased DGE analysis of RBPs. This revealed that acute endurance exercise in young individuals had a minimal impact on DGE of RBPs in comparison to the effect acute endurance exercise in muscles of older individuals (Figure 3).

Interestingly, older trained individuals had fewer differentially expressed RBPs in response to acute exercise compared to acute exercise in sedentary individuals (Figure 3). This finding is in line with recently published data that indicated lower RBP expression levels in masters athletes compared to age-matched non-athletes.^70,71^

Becoming or remaining physically active, as well as performing regular exercise, shifts to play a more important role in health as one ages than during their youth. Recent findings have indicated that AS becomes dysregulated with aging in several tissues, including skeletal muscle.^16,71,72^ When aged skeletal muscles were compared to younger muscles, proteins of the spliceosome and several RBPs were overrepresented, leading to a change in the expressed variants of mRNA.^16^ Interestingly, when octogenarian masters athletes were compared to age-matched non-athletes, the athletes had lower protein expression of the spliceosomal and several RBPs.^70^ The authors went on to hypothesize that the underrepresentation of the spliceosomal proteins was related to the higher functioning mitochondria and the lower need for energy to be expended for splice-isoform generation.^70^ Thus, further research is needed to link exercise, AS regulation, and mRNA variant generation to the adaptive process in skeletal muscle of both younger and older adults.

Next, we used rMATS to explore the type and number of AS events at the exon level from the SRS from humans (Figure 4). To our knowledge, rMATS has not previously been used to explore how exercise may modify AS, allowing us to make several novel insights. First, it is apparent that SE is the dominant form of AS event category in response to an acute bout of exercise while A5SS and A3SS are potentially subjected to other factors, such as age. The second most abundant type of splicing event across all data sets was RI (Figure 4). Both SE and RI events have important implications on the transcriptome and proteome that should be explored in future studies, but these data provide novel mechanistic insights into how acute endurance exercise is modifying gene expression and potentially skeletal muscle responses and adaptations to exercise. Second, we observed that there were fewer splicing events in the muscles of younger individuals compared to older individuals in response to exercise. However, we interpret this cautiously due to technical limitations and differences with how the samples were collected and the way the sequencing was performed between the data sets. Future studies will be needed to test this important question more carefully. A third observation is that older skeletal muscles utilized more A5SS and A3SS compared to younger muscles in response to a bout of endurance exercise. This could be due to the different responses of signal transduction pathways to modify the splicing machinery in older muscles compared to younger muscles. Alternatively, this could indicate greater energetic stress in older muscles leading to different signaling pathways being activated, resulting in the utilization of weaker splice sites. This could have important ramifications on the transcriptome and proteome following exercise in older muscles and should be explored in more detail in future research.^73,74^

While SRS is a powerful technology and our data above illustrate its ability to detect DGE and exon level splicing events, it lacks the ability to detect full-length transcripts, which have great implications on the expression of protein isoforms. Thus, we wanted to determine if additional insights could be made concerning how exercise mechanistically impacts skeletal muscle by using LRS. First, we were able to separate DGE and DAS using the LRS data (Figure 5A,B,C). This revealed that a significant portion of genes only undergo DAS, meaning previous post-exercise DGE analyses would not have detected these changes even though they would have likely impacted the post-exercise protein levels in muscle. Next, we analyzed isoform switching, specifically in genes that were not differentially expressed following exercise. We observed a robust isoform-switching response at all three time points in recovery from endurance exercise (Figure 5D). Further analysis illuminated a significant enrichment of RBPs that undergo isoform-switching events during recovery from endurance exercise (Figure 6A,B,C). In other words, RBPs and splicing factors undergo regulatory alteration by genes encoding these proteins via AS. Such regulatory process is referred to as either auto-regulation (whereby a RBP regulates the splicing of its own transcripts) or cross-regulation (whereby a RBP regulates the splicing of other RBP-coding transcripts). This idea of auto- and cross-regulation as a means of regulating functional levels of RBPs is not a novel concept in the field of RNA biology, as previously demonstrated in neuronal (NOVA1)^75^ and cancer (PTBP1/2 and SRSF3)^76^ cells.

Interestingly, several of the isoform-switching events lasted at least 24 hours into recovery from exercise, indicating a lasting (i.e., not transient) re-wiring of the transcriptome that likely impacts the proteome as we have shown is the case for hnRNP-A3 (Figure 6D–G).

Splicing and AS are regulated by a multitude of factors including (but not limited to) transcriptional rate, chromatin structure, the spliceosome, and RBPs.^4,5^ The potential for exercise to modify the activity of the spliceosome, as well as post-translational modification of RBPs which may impact binding to their respective target RNAs (and thus modify their splicing), is particularly important. RBPs bind to RNAs and impact their entire lifecycle: from splicing to localization, to translation, and degradation.^18^ Specifically, RBPs bind to *cis*-regulatory elements (i.e., intronic/exonic splicing enhancers and silencers) to regulate the inclusion of specific exons, resulting in the expression of different transcript variants and protein isoforms. Differential use of isoforms between conditions (isoform switching) can have profound biological impacts, even if they are only transiently induced by exercise, due to the difference in functions of the isoforms. Functional consequences of isoform switches can include gain or loss of protein domains or signal peptides, loss of protein coding sequence, and change in propensity for nonsense-mediated decay.^13^ Isoform switches are implicated in many diseases, and it has already been observed following exercise (PGC-1α)^15^; however, the full extent and impact of isoform switching post-exercise is far from understood.

It is highly likely, but not yet tested, that exercise remodels the binding patterns of RBPs either through post-translational modifications of the RBPs themselves or through isoform switching and AS changes to RBPs themselves, which would modify the target genes of RBPs. Metabolic stress responsive pathways, such as AMPK, are already known to regulate post-translation modifications of RBPs.^21^ Interestingly, many RBPs tend to regulate their protein abundance through AS and isoform switching. Our isoform switching data indicate an enrichment of events in RBPs post-exercise. This indicates that exercise may remodel the expression of protein isoforms by modifying the function of RBPs. Our findings not only support the preceding study from our laboratory that focused on SF3B4 and SRSF2,^30^ but others that have observed similar phenomenon in a separate setting: Henrich et al.^77^ illustrated 9 weeks of microgravity conditions leading to AS-driven diversification of the transcriptome in mouse skeletal muscle, including RBPs (e.g., *mMbnl1* and *mRbfox1*). This mechanism is important in shaping the transcriptome and proteome in response to specific cellular stresses. Bringing this idea to the study of skeletal muscle adaptations in response to exercise is an innovative and novel molecular mechanism of adaptation (Figure 7).

**Figure 7.**
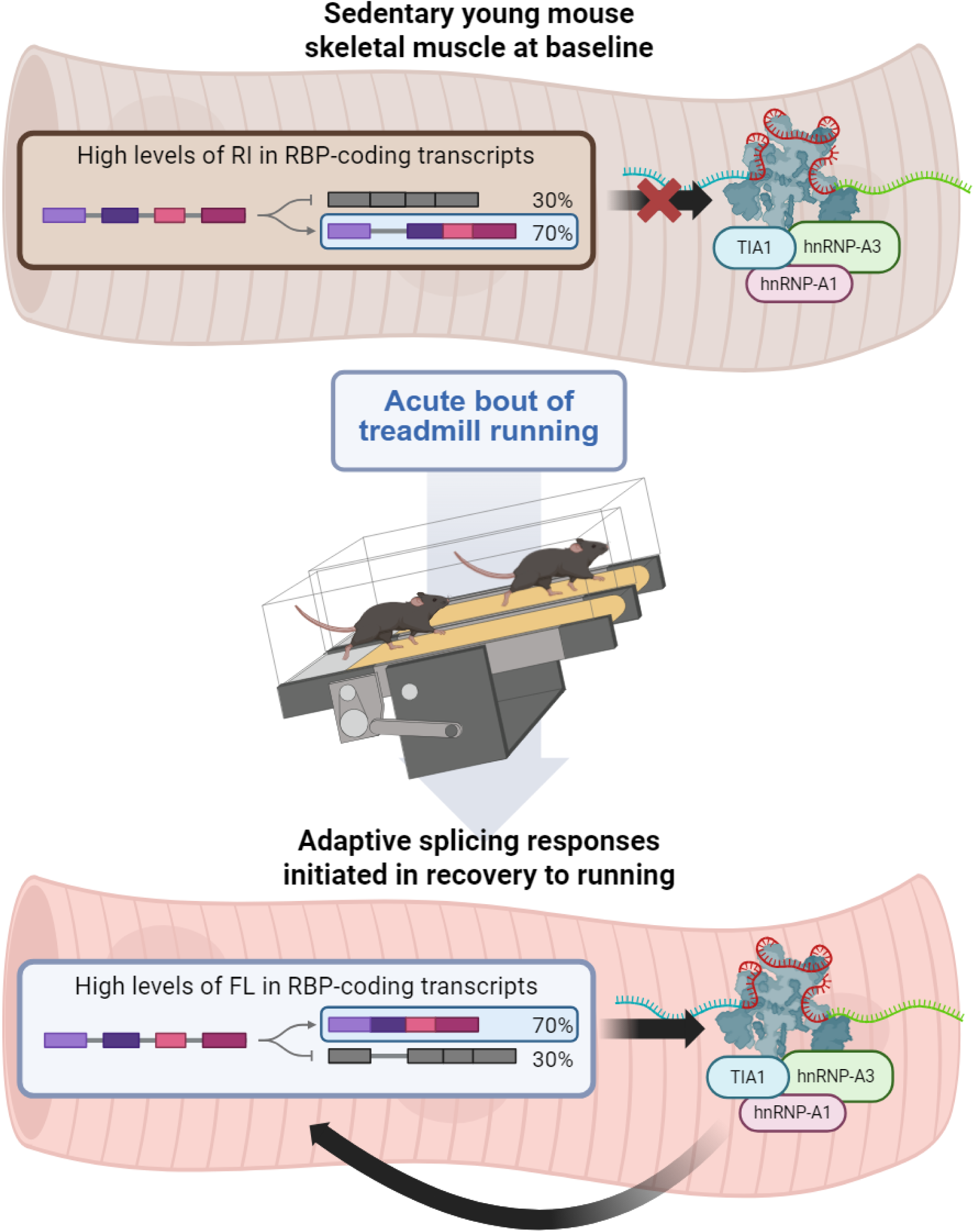
Current model on the effects of acute endurance exercise on the AS regulatory mechanism with key *trans*-factors/RBPs (e.g., hnRNP-A1, -A3, and TIA1).

The present study is not without limitations. The human SRS data utilized for this study included on average 20 million reads per sample, which is insufficient/minimum threshold to yield biologically true isoform-switching analyses with IsoformSwitchAnalyzeR. Hence, there is a growing need for employing LRS technologies to better examine alternative splicing and isoform switching in the field of exercise biology. Additionally, the mice cohort had mixed sexes unlike the human cohorts utilized for analyses, which may further limit the comparability. Indeed, protein analysis was only conducted for one of the RBPs, hnRNP-A3, but not at the global level for this study. To better validate the significance of DAS and isoform-switching events, future studies will incorporate proteomics data with LRS data via proteogenomic approaches.

In summary, the present data support recent evidence of divergence in the skeletal muscle AS-related responses and RBP expression by age and training status. Moreover, LRS is the preferrable method of transcriptomics study when exploring AS and isoform-switching events, whereas utilizing SRS may require cost prohibitive read depth per sample for many groups (at least 100 million reads per sample) to proficiently conduct the isoform switching analyses.

Additionally, our acute endurance exercise mouse model demonstrated that, even without any changes in overall gene expression, the protein expression of hnRNP-A3 was increased by nearly two-fold in response to exercise for at least 24 hours via isoform switching with the power of LRS. Therefore, our study points out that AS following exercise in skeletal muscle is an underexplored and untapped area, ripe for future studies to increase our mechanistic understanding of exercise.

## Supporting information

All supplemental figures

## IV: DATA AVAILABILITY STATEMENT

The human SRS data were extracted from deposits by Pattamaprapanont et al.^31^ and Rubenstein et al.^32^ in the NCBI Gene Expression Omnibus database with the accession numbers GSE87749 and GSE151066, respectively. The mouse LRS data are deposited in the NCBI Gene Expression Omnibus database with the accession number GSE279359. All data used to generate the figures in this manuscript will be deposited at the Deep Blue Data repository with a permanent DOI (DOI will be provided upon publication).

## V: CONFLICT OF INTEREST STATEMENT

The authors declare that the research was conducted in the absence of any commercial or financial relationships that could be construed as a potential conflict of interest. The authors declare no competing interests.

## VI: AUTHOR CONTRIBUTIONS

AA, ALS, JJK, and ATL conceived and/or designed the research. AA, ALS, JJK performed the research and acquired the data, AA, JJK, JYY, EYZ, and ATL analyzed and interpreted the data. All authors were involved in drafting and revising the manuscript.

## VII: ACKNOWLEDGMENTS

This research was supported by the National Institutes of Health and National Cancer Institute under award number 5R00CA197672-04, Michigan Integrative Musculoskeletal Health Core Center (MiMHC) P30 grant from NIH/NIAMS under award number P30 AR069620, and the University of Michigan’s School of Kinesiology Marie Hartwig Research Fund.

## Abbreviations

A3SS: alternative 3’ splice site
A5SS: alternative 5’ splice site
AS: alternative (RNA) splicing
ATSS: alternative transcription start site
ATTS: alternative transcription termination site
DAS: differential alternative splicing
DGE: differential gene expression
DIE: differential isoform expression
hsAA: *homo sapiens* aged active
hsAS: *homo sapiens* aged sedentary
hsYS: *homo sapiens* young sedentary
LRS: long-read RNA sequencing
mmYS: *mus musculus* young sedentary
MXE: mutually exclusive exons
RBP: RNA-binding protein
RI: retained intron
SE: skipped exon
SRS: short-read RNA sequencing.

## Notes

### Competing Interest Statement

The authors have declared no competing interest.

### Summary of Updates

A significant revision to the data analysis and all figures and data was made.

