## Supplementary figures and images for "Impact of Acute Endurance Exercise on Alternative Splicing in Skeletal Muscle"

### All supplemental figures

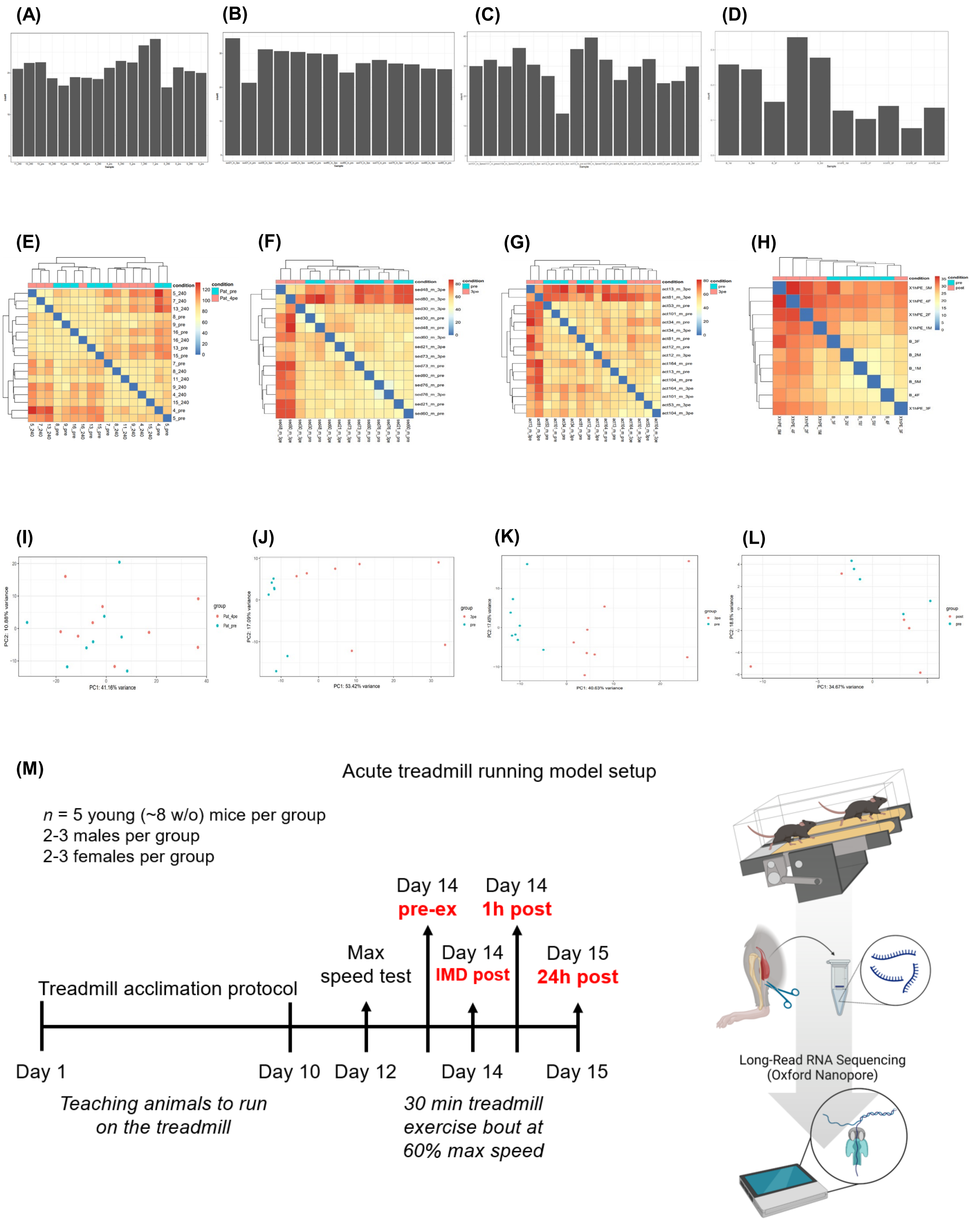

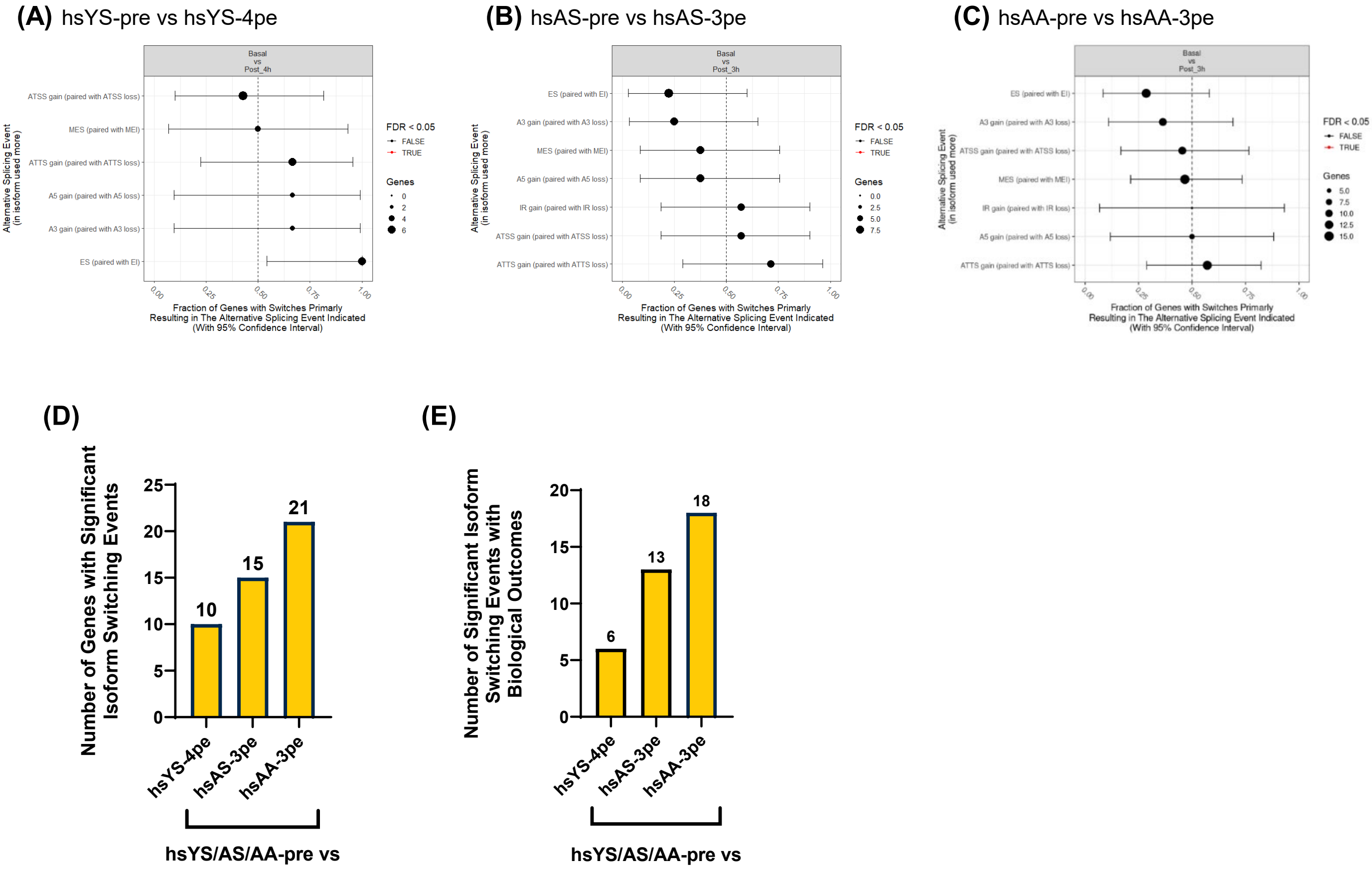

#### Figure S3.

**(A)** hsAS-pre:3pe rMATS

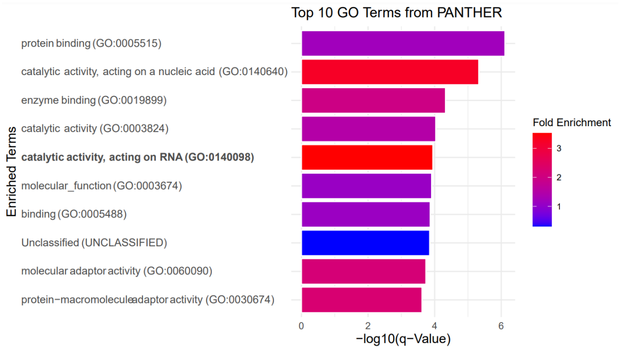


**(B)** hsAA-pre:3pe rMATS

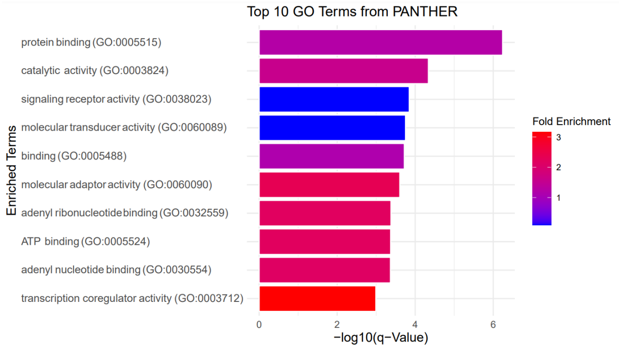

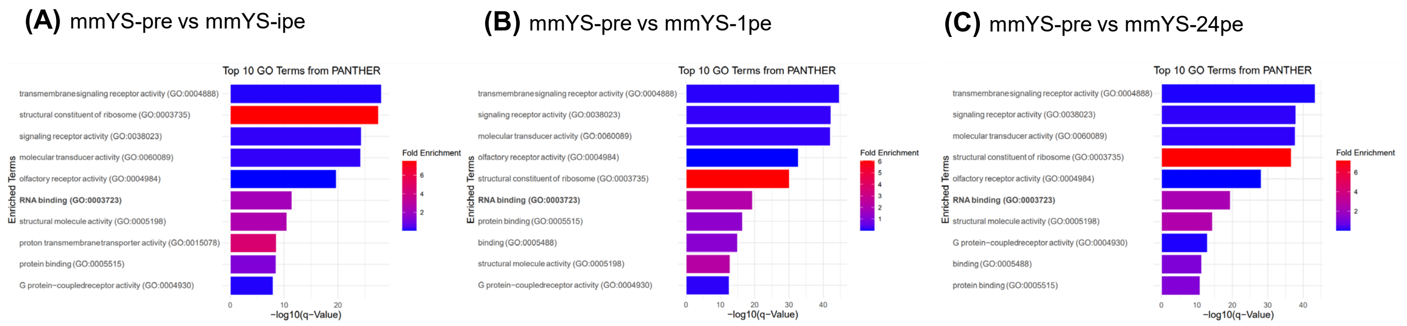

**Figure S5.**

**(A)** *m18S* rRNA

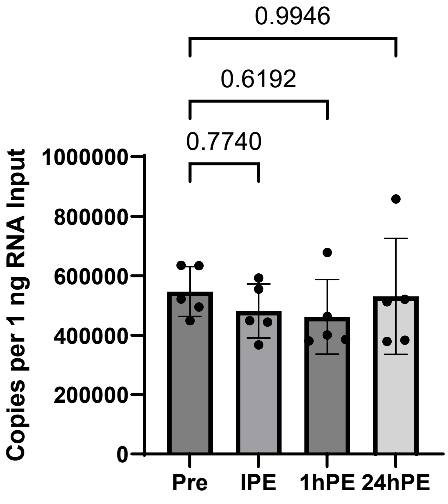


**(B)** *mHnrnpa1* Gel-Based RT-PCR

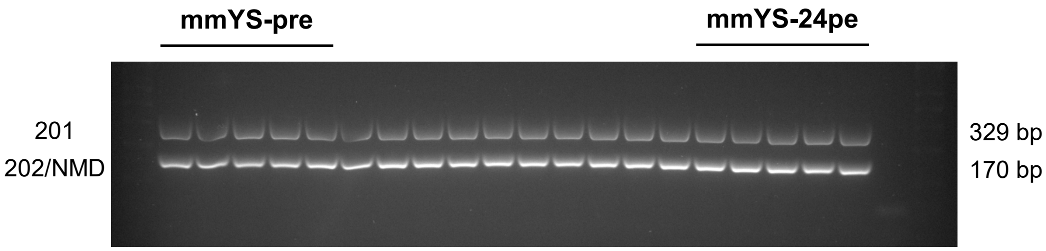

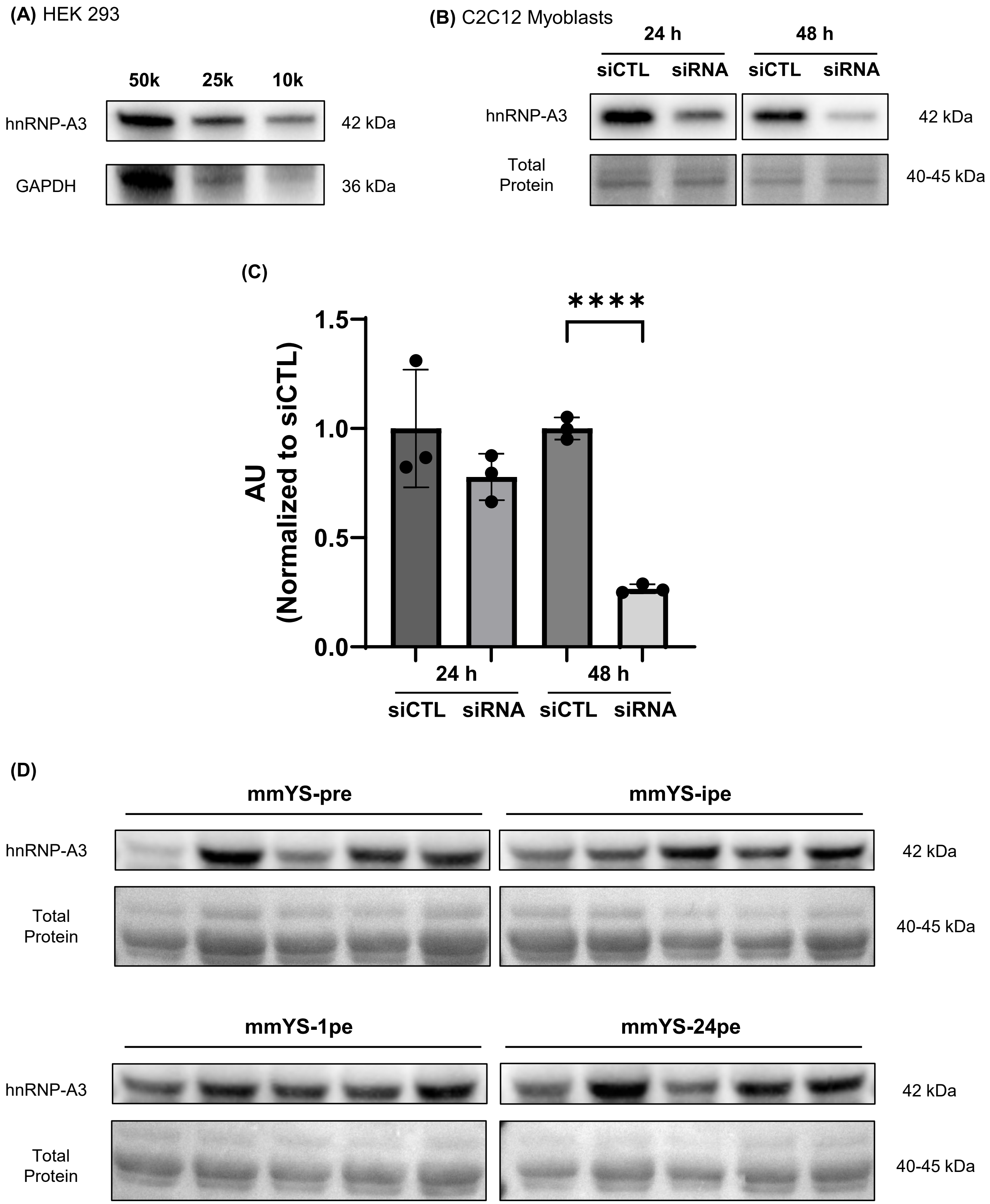
