## Supplemental figure legends for "Impact of Acute Endurance Exercise on Alternative Splicing in Skeletal Muscle"

**Figure S1.** RNA-Seq data quality metrics and animal acute treadmill running model. Bar graph of total read count of (A) hsYS, (B) hsAS, (C) hsAA, and (D) mmYS cohort data by sample in millions. Euclidean hierarchical heatmap analysis of (E) hsYS, (F) hsAS, (G) hsAA, and (H) mmYS cohort data by sample. Principal component analysis of (I) hsYS, (J) hsAS, (K) hsAA, and (L) mmYS cohort data by sample. (M) Overview of treadmill acclimation protocol, max speed test, and pre-/post-acute exercise time points of skeletal muscle harvests.

**Figure S2.** Forest plots of the differences in the faction of genes stratified in AS event type across (A) hsYS, (B) hsAS, and (C) hsAA cohorts compared between pre- and post-acute bout of exercise using IsoformSwitchAnalyzeR. (D) Isoform switching genes with statistical significance (ΔΠ > 10%, *p*-adj < .05) between pre- and post-acute exercise time points across the human cohorts. (E) Isoform switching genes with statistical significance and predicted biological outcomes between pre- and post-acute exercise time points across the human cohorts.

**Figure S3.** Differential splicing events of hsAS and hsAA. (A) Top 10 GO terms of enriched genes with splicing events of hsAS-pre and hsAS-3pe using PANTHER. (B) Top 10 GO terms of enriched genes with splicing events of hsAA-pre and hsAA-3pe using PANTHER.

**Figure S4.** Significant isoform-switching genes of mmYS in the skeletal muscle comparing pre- and post-acute exercise using Swan. (A) Top 10 GO terms with significant isoform-switching events of mmYS-pre and mmYS-ipe using PANTHER. (B) Top 10 GO terms with significant isoform-switching events of mmYS-pre and mmYS-1pe using PANTHER. (C) Top 10 GO terms with significant isoform-switching events of mmYS-pre and mmYS-24pe using PANTHER.

**Figure S5.** (A) RT-ddPCR of *m18S* rRNA illustrated equal loading of the mouse cDNA samples for gel-based RT-PCR and RT-ddPCR. (B) Full image of gel-based RT-PCR of *mHnrnpa1* found in Figure 6I.

**Figure S6.** Western blot validation of anti-hnRNP-A3 antibody in human and mouse samples. (A) Wild-type HEK 293 cells were used to verify cross-species specificity of the hnRNP-A3 antibody and was loaded by 50k, 25k, and 10k cell equivalents. Loading determined with GAPDH. (B, C) C2C12 myoblasts were transfected with 30 nM of either siCTL or *Hnrnpa3* siRNA for 24 and 48 h to validate protein specificity of the antibody. Total protein with Pierce™ Reversible Protein Stain Kit. (D) Full collection of western blots used to analyze significant isoform switching of hnRNP-A3 at the protein level, normalized to total protein (Ponceau S stain) at 40-45 kDa band detection then normalized to mmYS-pre for relative quantification in AU. ****, *p* < 0.0001 compared to siCTL.
